# FACT-mediated maintenance of chromatin integrity during transcription is essential for mammalian stem cell viability

**DOI:** 10.1101/2021.06.21.449309

**Authors:** Imon Goswami, Poorva Sandlesh, Aimee Stablewski, Ilya Toshkov, Alfiya F Safina, Mikhail Magnitov, Jianmin Wang, Katerina Gurova

**Author notes:** Corresponding authors: Katerina Gurova, Department of Cell Stress Biology, Roswell Park Comprehensive Cancer Center, Elm and Carlton Streets, Buffalo, NY, USA, 14263., Jianmin Wang, Department of Biostatistics and Bioinformatics, Roswell Park Comprehensive Cancer Center, Buffalo, NY, USA, 14263. Department of Neurological Surgery, University of Pittsburgh, Pittsburgh, PA University of Pittsburgh, Hillman Cancer Institute, Pittsburgh, PA. These two authors contributed equally to the work (Imon Goswami, Poorva Sandlesh).

## Abstract

Preservation of nucleosomes during replication has been extensively studied, while the maintenance of nucleosomes during transcription has gotten less attention. The histone chaperone FACT is involved in transcription elongation, although whether it disassembles or assembles nucleosomes during this process is still unclear. We deleted the FACT subunit in adult mice to clarify the function of FACT in mammals. FACT loss was lethal due to the loss of the earliest progenitors in bone marrow and intestine, while mor differentiated cells were not affected. Using cells isolated from several tissues, we showed that FACT loss was lethal only for stem cells but not cells differentiated *in vitro*. FACT depletion led to increased chromatin accessibility in a transcription-dependent manner, suggesting that nucleosomes are lost during transcription in the absence of FACT. The most prominent response to the loss of nucleosomes was the activation of interferon signaling and the accumulation of immunocytes in sensitive organs. FACT maintained chromatin integrity during transcription in mammalian adult stem cells, suggesting that chromatin transcription in these cells is different from more differentiated cells.

## INTRODUCTION

Chromatin organization defines cell identity by limiting the access of transcription machinery to DNA in a region-specific manner, making the existence of cell- and condition-specific transcriptional programs possible. Although significant progress has been made in elucidating the structure and mechanism of action of factors involved in chromatin maintenance, studies have mainly used cell-free conditions or simplified model organisms (e.g., yeast or human tumor cells). The role of these factors in modifying gene expression in a cell-specific manner in multicellular organisms, including the spatio-temporal regulation of their activities, is still far behind.

One group of such factors is the histone chaperones that have long been considered a component of the basic replication and transcription machinery responsible for the assembly and disassembly of nucleosomes [1]. However, studies done in multicellular organisms demonstrated that they are not ubiquitously expressed and that their depletion has different consequences for different cells [2–5]. These characteristics suggest that either their function is not unique or the control of chromatin integrity may not be required in all cells.

Histone chaperone FACT (FAcilitates Chromatin Transcription) is especially interesting in this regard. In recent years, the view of its function as an essential transcription elongation factor facilitating the access of the transcriptional machinery to nucleosomal DNA has been questioned by several mechanistic and structural studies [6], [7]. FACT does not bind to nucleosomes fully wrapped with DNA but binds to nucleosomes with partially unwrapped DNA [8–10]. Reduced FACT levels in several cell types increase chromatin accessibility at different genomic regions and are accompanied by elevated transcription [11, 12], suggesting that FACT may restrict the access of the transcriptional machinery to DNA. The phenotypes of FACT knockout in *C. elegans* or *D. rerio* do not support the proposed function of FACT as an essential transcription elongation factor [13, 14]. Different laboratories have demonstrated that several normal mammalian cell types do not depend on FACT for growth or viability [11, 14, 15]; however, most mouse and human tumor cells cannot grow or die after FACT knockdown [12, 16]. It is still not clear what makes tumor cells dependent on FACT. This knowledge could be valuable for cancer research. The absence of FACT expression in most adult mammalian cells [17] suggests that FACT may have cell-specific functions or that some, but not all, cells require FACT due to a specific chromatin state. To understand FACT function in multicellular eukaryotes, it is critical to identify which mammalian cells depend on FACT and which processes are disturbed in these cells upon FACT loss.

FACT consists of two subunits, SPT16 (Suppressor of Ty 16) and SSRP1 (Structure-Specific Recognition Protein 1). General knockout (KO) of the *Ssrp1* gene in mice is lethal at the blastocyst stage [18] when expression of the FACT subunits is the highest in the life span of mice [17]. However, whether FACT is essential for the viability of adult mice when its expression is very low is not known. The stability of both FACT subunits depends on their binding to each other, and depletion of one leads to the disappearance of the other [19]. Based on this knowledge, we generated an *Ssrp1* conditional knockout (KO) mouse model to eliminate FACT and observe cell-specific and age-dependent consequences of FACT loss. Specifically, we crossed mice with LoxP sites inserted into the introns of the *Ssrp1* gene and R26-CreER^T2^ mice to achieve FACT depletion in adult animals using tamoxifen administration [20]. In this study, we determined the phenotypes of FACT loss in mice at different ages and identified the FACT-dependent cells in FACT-sensitive organs. We also propose the reasons for this selective FACT dependency.

## Results

### 1. FACT loss in mice is lethal at all ages

For this study, we compared mice homozygous for the *Ssrp1* mutant allele, *Ssrp1^fl/fl^,* with heterozygous and wild-type animals [20]. Consistent with a previous report [18] we demonstrated that mice with one functional *Ssrp1* allele lacked any phenotype. Therefore, we used these heterozygous mice as controls in many experiments. These mice were crossed with Rosa26 CreERT2 (*CreER^T2+/+^*) mice to obtain mice homozygous for both alleles since the presence of one copy of *CreER^T2^* results in poor or no excision of *Ssrp1* [20]. Treatment of the mutant mice with tamoxifen for five days led to the reduction and disappearance of both FACT subunits in the spleen, the only organ in adult mice where SSRP1 and SPT16 can be detected by immunoblotting (Fig.EV1A). Based on this finding, tamoxifen was administered for five days in subsequent experiments.

One week after the end of tamoxifen administration, both female and male mice began losing weight and died or were euthanized between 12 and 50 days after the end of treatment (Fig.1A, B). Because Cre can cause DNA damage due to its non-specific recombinase activity, we tested the effect of tamoxifen administration on mice with two CreER^T2^ alleles and wild-type *Ssrp1* and did not observe any toxicity (Fig.EV1B), indicating that FACT loss caused the deaths of *Ssrp1^fl/fl^* mice. Older male mice (24–40 weeks at the start of tamoxifen administration) lived longer than younger males or female mice; however, they still died after FACT loss (Fig.1B). A comparison of the complete blood counts (CBC) between male and female *Ssrp1^fl/fl^; CreER^T2+/+^* mice treated with tamoxifen or vehicle control showed a significantly reduced number of white blood cells (i.e., lymphocytes and granulocytes) but no changes in the numbers of red blood cells and platelets (Fig.1C). Serum biochemistry analysis did not find a significant deviation between tamoxifen treated *Ssrp1^fl/fl^; CreER^T2+/+^ and Ssrp1^fl/+^; CreER^T2+/-^* mice except slight reduction of lipase in homozygous mice (Table EV1). Tamoxifen-treated *Ssrp1^fl/fl^; CreER^T2+/+^* mice showed signs of dehydration, and some animals developed atopic dermatitis-like skin lesions (Fig.1D). Subcutaneous fluids (1-2 mL/day) did not rescue these mice (Fig.EV1C). Thus, although FACT is only present in a limited number of cells in several organs of adult mice [17], it is essential for the viability of adult mice. For subsequent experiments, we used mice between 10 and 15 weeks of age.

**Figure 1.**
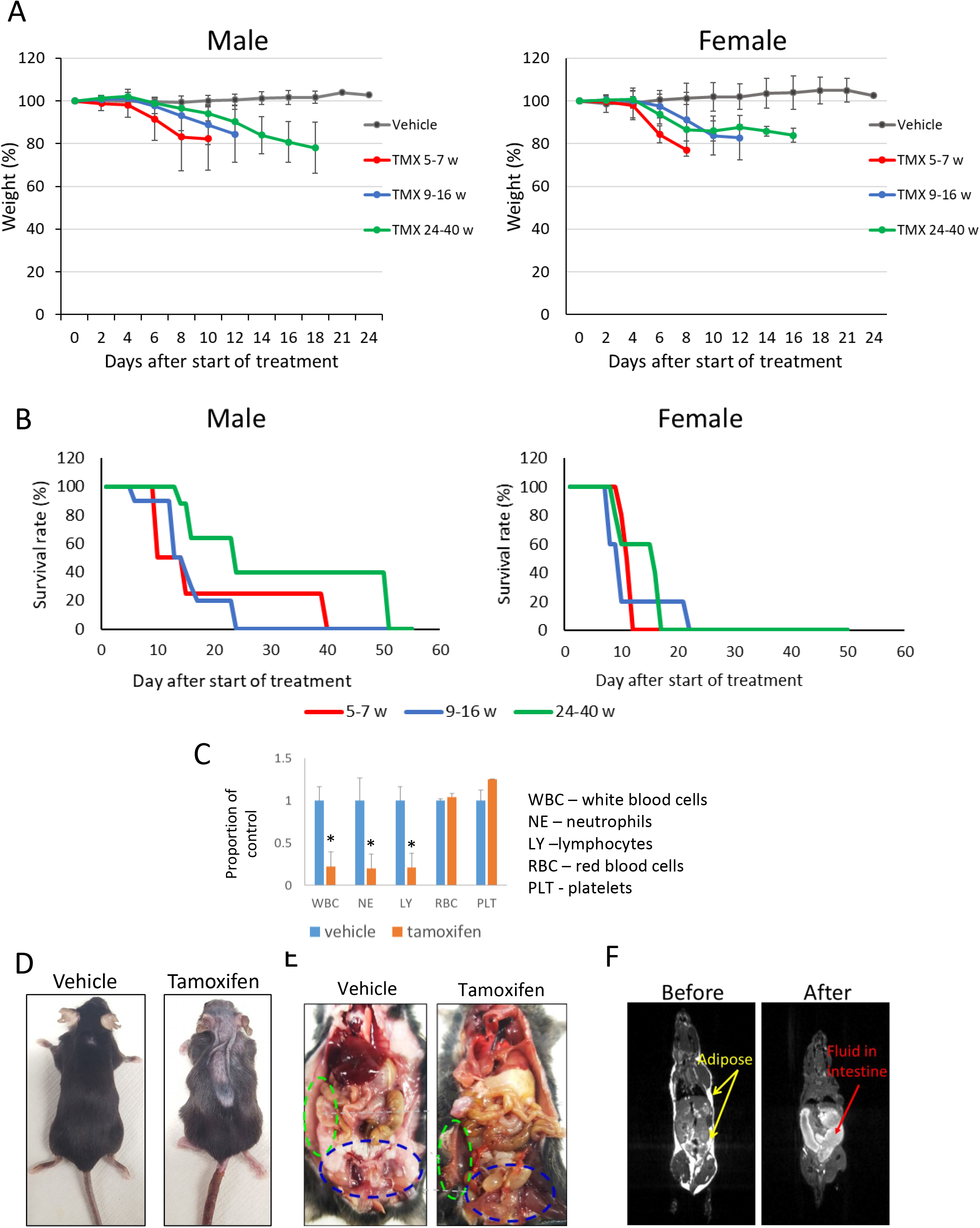
Effect of *Ssrp1* KO in adult mice. A, B. Normalized weight (A) and survival (B) of *Ssrp1^fl/fl^; CreER^T2+/+^* mice of different ages (weeks) after the start of tamoxifen (TMX) treatment. A. Mean weight (%) ± SD. P < 0.05 for all tamoxifen treated groups versus vehicle (ANOVA). B. Kaplan-Meier survival curves. Log-rank test p = 0.032 for the oldest male age group versus all other male groups; p = 0.013 for all females versus all males; for all other comparisons p > 0.05. *n = 6-10 mice*. C. Normalized blood cell counts in *Ssrp1^fl/fl^; CreER^T2+/+^* mice treated with vehicle or tamoxifen on day 7 after start of treatment. Mean ± SD, * indicates p < 0.05 (unpaired t- test). *n = 3 mice*. D. Photographs of vehicle- or tamoxifen-treated *Ssrp1^fl/fl^; CreER^T2+/+^* mice seven days after start of treatment. E. Gross pathology examination of *Ssrp1^fl/fl^; CreER^T2+/+^* mouse euthanized due to 20% weight loss after treatment with tamoxifen. Vehicle-treated mouse is shown for comparison. Green and blue ovals indicate subcutaneous and inguinal fat deposits, respectively. F. MRI image of *Ssrp1^fl/fl^; CreER^T2+/+^* mouse before and after tamoxifen treatment.

### 2. Several tissues are sensitive to FACT depletion

Gross pathology examination revealed complete loss of adipose tissue (Fig. 1E, F) and accumulation of liquid in the large intestine (Fig.1E) following the loss of FACT. The weights of the major parenchymal organs were not significantly changed, except the spleen, which was smaller in the tamoxifen-treated groups (Fig. EV1E). The ovaries were also significantly reduced in the tamoxifen-treated mice (Fig.EV1F); however, these changes have been observed in mice with wild-type *Ssrp1* and reported in the literature as a direct effect of tamoxifen treatment [21]. In contrast, the testes, which also have high FACT expression [17], were not reduced by tamoxifen treatment (Fig.EV1E, G).

Upon FACT depletion histological changes were found in organs that express FACT subunits in basal conditions [17]. Reduced cellularity was observed in the bone marrow and spleen, including atrophic changes in the red pulp (Fig.2A), areas where FACT is present in adult mice [17]. High FACT levels are also present in cells at the bottom of the crypts of the small and large intestines [17]. *Ssrp1* KO resulted in the thinning of the colon villi and the emergence of a small number of degenerative crypts (Fig. 2A, B). No histological changes were seen in the high FACT- expressing testes (Fig. EV1G). There was also no reduction in FACT expression in the testes after tamoxifen administration (Fig. EV2A), suggesting that recombination did not occur in the testes, most probably due to the limited penetration of tamoxifen through the blood-testis barrier [22]. Among the FACT-negative organs, tamoxifen administration causes histological changes only in the liver; there was a loss of glycogen granules in the hepatocytes. Some of the hepatocytes were enlarged or undergoing mitosis, which is not typical for the adult liver. Lymphoid cell infiltration was also seen in some areas (Fig.2C). The skin was excluded from the histological analysis in this study because the skin lesions first appeared in mice when they lost almost 20% of their weight, requiring euthanasia. Accurate assessment of the consequences of FACT loss in the skin will require mouse models with keratinocyte-specific expression of CreER^T2^.

**Figure 2.**
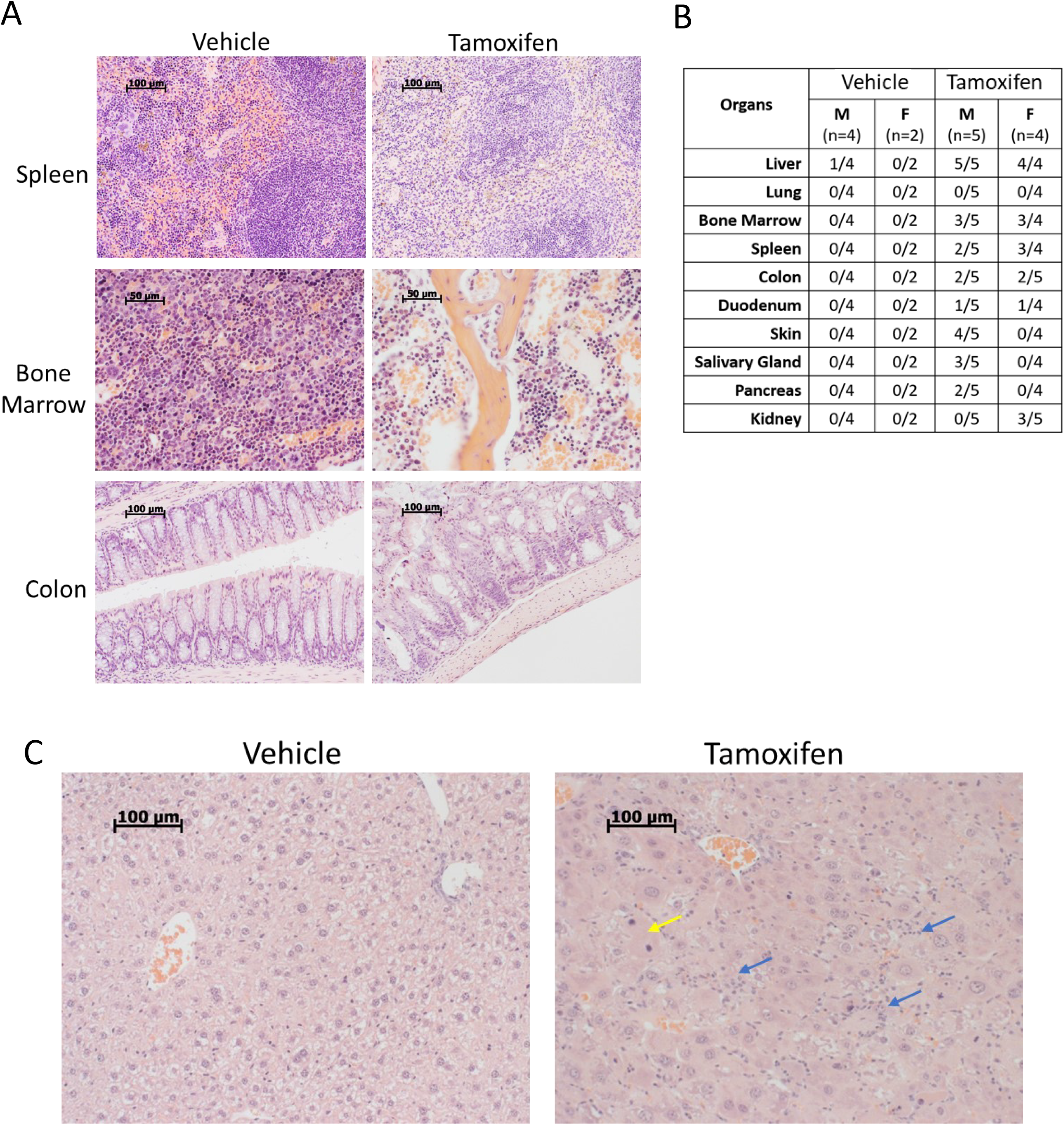
Pathological findings in *Ssrp1^fl/fl^; CreER^T2+/+^* mice after tamoxifen treatment. A. H&E staining of mouse organ sections. Scale bars: spleen, colon 100 μM, bone marrow – 50 μM. B. Incidence of pathological findings (number of mice with pathology/total number of mice). C. H&E staining of liver sections. Yellow arrow - a mitotic hepatocyte; blue arrows - lymphocyte infiltration. Scale bars – 100 μM.

Altogether these findings may indicate the suppression of hematopoiesis and the suppression of the renewal capacity of the intestine, leading to atrophic changes that could explain the observed dehydration because the large intestine is the major organ of water absorption. Changes in the liver and reduction of lipase concentration in serum are likely secondary to weight loss. The loss of adipose tissue could explain the weight loss and might be either primary (death of adipocytes) or secondary (loss of fat from cells due to malnutrition and dehydration) to FACT loss.

### 3. FACT loss impairs proliferation and induces cell death in several organs

To better understand the consequences of FACT depletion we compared the levels of proliferation and cell death in the major parenchymal organs between vehicle- and tamoxifen-treated mice using EdU labeling as a marker of DNA replication and cleaved caspase-3 staining as a marker of apoptosis. Three days after the end of tamoxifen treatment, the number of EdU-positive cells was significantly reduced in the spleen and intestines of *Ssrp1^fl/fl^; CreER^T2+/+^* mice (Fig.3A–C). Although EdU-positive cells are normally absent from the liver, some EDU-positive cells did appear following tamoxifen treatment, consistent with the appearance of mitotic cells in the H&E-stained liver sections (Fig.2C and EV2B). Changes in EdU positive cells in the small and large intestines were moderate, with some sections with no changes in the number of EdU-positive cells and some areas with a reduced frequency of replicating cells in the lower portion and the spread of replicating cells towards the upper portion of the crypts (Fig.3B, C). In the spleen, intestines, and colon, we observed a slight increase in the number of cleaved caspase-3- positive cells; however, they were not concentrated in any specific cellular zone in these organs (Fig.3D and EV2C). Thus, FACT depletion reduced the number of cells in the hematological organs and intestines. This loss of cells was predominantly due to reduced cell proliferation of cells. Interestingly, hepatocytes started proliferating in the liver upon *Ssrp1* KO. Thus, a general impairment of DNA replication or mitosis, processes involving FACT activity [23–27], cannot explain the toxicity of *Ssrp1* KO.

**Figure 3.**
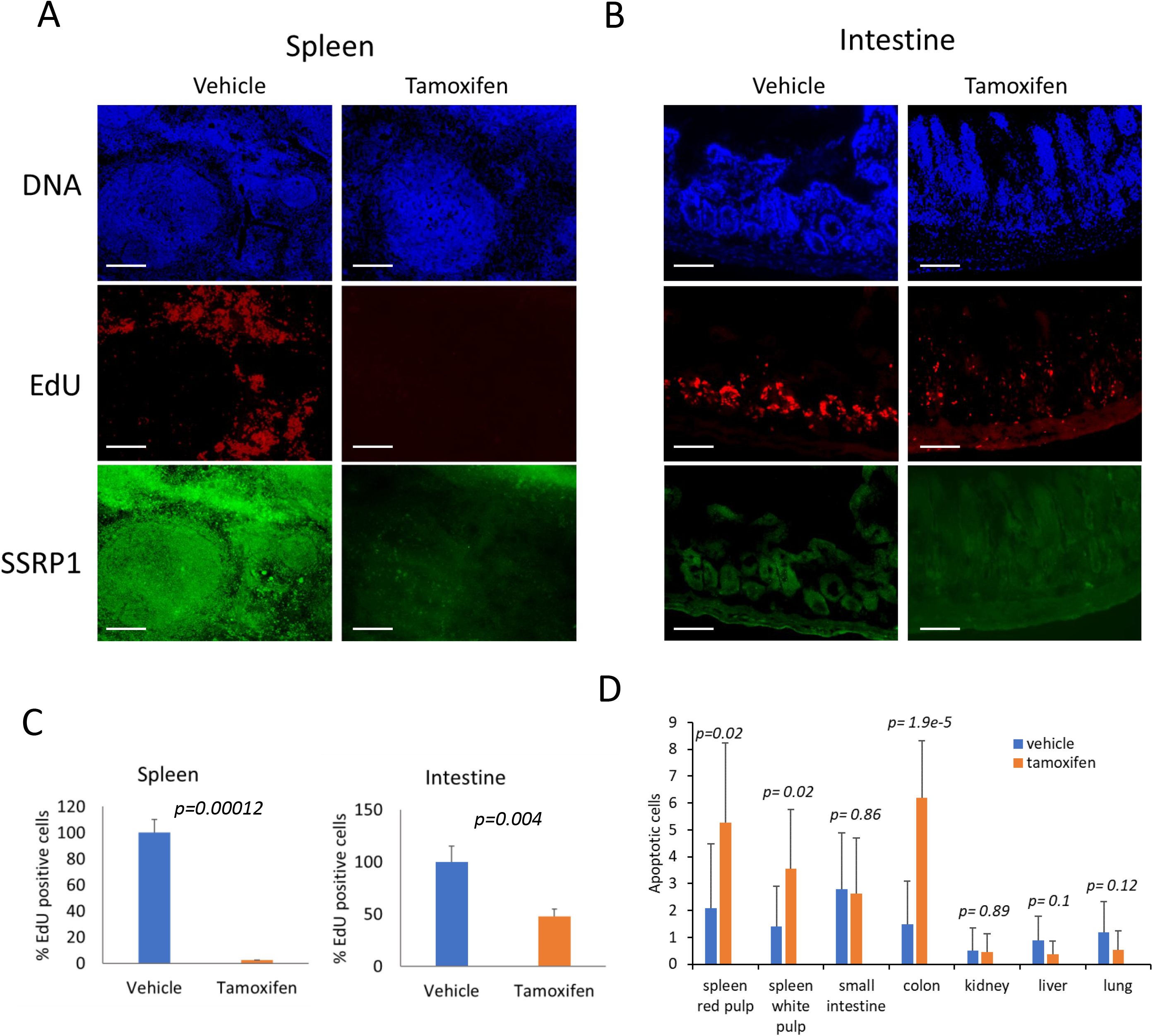
FACT depletion in mice leads to reduced proliferation and cells death in several organs. A, B. Effect of tamoxifen on replication in the spleen (A) and intestine (B) of *Ssrp1^fl/fl^; CreER^T2+/+^* mice. EdU was given to mice three days after the end of tamoxifen treatment (1 hour before tissue collection). Organ sections were stained for total DNA (Hoechst), EdU, and SSRP1. Scale bars – 100 μM. C. Quantitation of EdU staining. Mean number of EdU positive cells per field of view +/- SD. *n ≥ 5 fields of view*. 3 mice were analyzed. D. Quantitation of cleaved caspase-3 staining. Mean number of positive cells per fields of view +/- SD, *n ≥ 5 fields of view.* Two male and female mice were analyzed. P-values for C and D were calculated using the unpaired t-test.

### 4. FACT depletion is accompanied by the loss of long-term hematopoietic stem cells from bone marrow and Lgr5 positive stem cells from the intestines

To identify cells sensitive to FACT loss in hematopoietic and intestinal tissues, we performed single-cell RNA sequencing (scRNA-seq). Quality control confirmed the integrity of the samples with viabilities of > 90% and > 75% for the cells obtained from the bone marrow and intestines respectively. There were a comparable number of cells sequenced (about 8,000–10,000 per sample), the number of reads generated (about 20,000–27,000), and the number of genes quantified (approximately 1000 per cell) (Fig. EV3A-B).

The Uniform Manifold Approximation and Projection (UMAP) plots for vehicle- and tamoxifen-treated samples showed significant overlap, suggesting that, in general, the tissue architecture was well preserved in both organs (Fig. EV3C). There were no significant changes in the distribution of cells at the different phases of the cell cycle within the clusters between the vehicle- and tamoxifen-treated samples and no preferential depletion of cells in any particular phase of the cell cycle (Fig. EV3D, E), suggesting that FACT loss did not block any specific phase.

A comparison of the bone marrow UMAP plots revealed several clusters with visibly changed numbers of cells. Three clusters (14, 15, and 16) almost completely disappeared after FACT depletion (Fig.4A). Using the Immunological Genome Project (ImmGen) database [28] as a reference, we identified the major types of bone marrow cells in our samples (Fig.4B). There were no dramatic differences in the proportions of the different cell types between samples from the vehicle- and tamoxifen-treated mice (Fig. 4C). In several instances, the clusters with increased or decreased numbers of cells belonged to the same cell type (e.g., the disappearing clusters 14–16 and the increasing cluster 9 were all identified as ‘stem cells’) (Fig.4D). The same was true for the neutrophils and B cells. Several clusters consisting of one cell type (cluster 11 – dendritic cells (DC), 20 – basophils (Bph), 21 – T cells, and 22 – NK cells) were slightly increased in number (Fig. 4C, D). Therefore, we examined the differences between the decreased and increased clusters of the same cell type.

**Figure 4.**
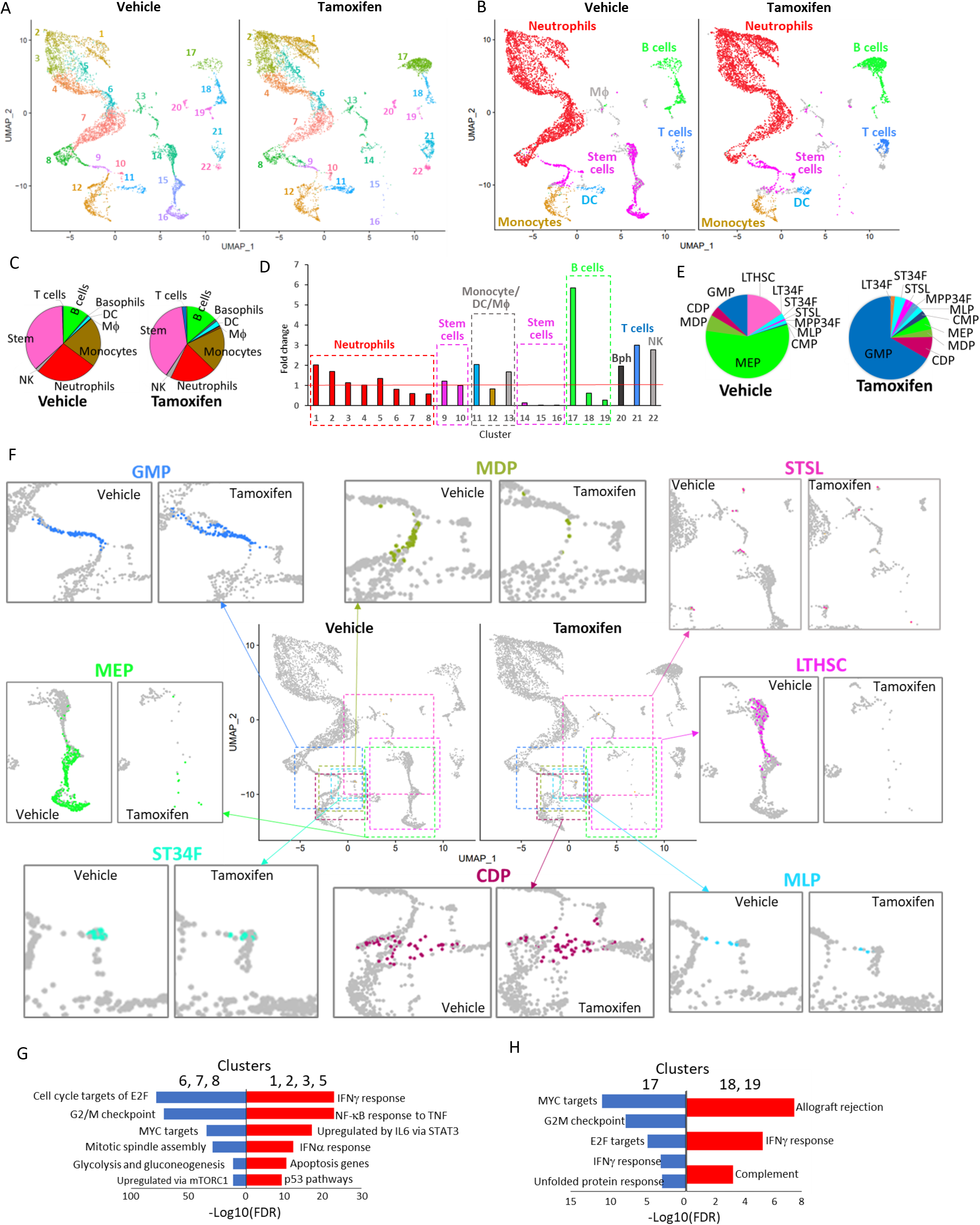
Effect of FACT depletion on cell composition of bone marrow from *Ssrp1^fl/fl^; CreER^T2+/+^* mice (pooled data from two mice on day 1 after the end of treatment). A. UMAP plots with unbiasedly identified clusters shown in different colors. B. UMAP plots colored according to cell type identified by transcriptional signatures using the Immgen database as a reference. C. Distribution of bone marrow cell types in vehicle- and tamoxifen-treated mice. The pie charts are colored accordingly to panel B. Only cell types with more than ten cells in any of the conditions are included. D. Fold-change in the proportion of cells in each cluster shown in panel A in samples from tamoxifen-treated versus vehicle-treated mice. Cell types are identified by color as in panel B. E. Distribution of different types of bone marrow stem cells from vehicle- and tamoxifen- treated mice. Stem cell types were identified by transcriptional signatures using the Immgen stem cell database as a reference. Only cell types with more than five cells in any of the conditions identified are presented. LTHSC – long-term hematopoietic stem cells (HSC), ST34F – CD34+Flk2- short-term (ST) HSC, STSL – ST reconstituting stem cells, MPP34F – CD34+Flk2+ multipotent progenitors, MLP – multi-lymphoid progenitors, CMP.DR – common myeloid progenitor, MEP – megakaryocyte-erythroid progenitors, MDP – monocyte-dendritic cell progenitors, CDP – common dendritic cell progenitors, GMP – granulocyte-monocyte progenitor. F. Changes in the abundance and distribution of major bone marrow stem cell types. Center – UMAP plots showing enlarged regions as colored dashed rectangles. The color of a rectangle corresponds to the color of the stem cells in individual sections. Around central UMAPs – enlarged sections of the central UMAP plot showing individual stem cell subtypes. Colored dots – corresponding stem cells, grey dots – other cells. G–H. GSEA of gene markers of neutrophil (G) or B cell (H) in clusters enlarged upon FACT loss (red) or reduced upon FACT loss (blue).

In the control samples, stem cells formed two separate groups: a central group isolated from other cells (clusters 14–16) and a group connecting the monocytes and granulocytes (clusters 9, 10) (Fig.4B). The central group of stem cells disappeared, while the cluster connecting the granulocytes and monocytes slightly increased (Fig.4B). We used the ImmGen datasets specific for bone marrow stem cells to determine the differences between these subpopulations. We identified 11 types of stem cells in our dataset (Fig.4E). The most dramatic change was observed for the long-term hematopoietic stem cells (LTHSCs), which completely disappeared after *Ssrp1* deletion (Fig.4E, F). CD34^-^ LTHSCs gave rise to CD34^+^ LTHSCs (LT34Fs), which increased in number; however, the total cell count was very low and unreliable (one cell in the control and four cells after FACT loss). The LT34Fs were followed by short-term HSCs (ST-HSCs), consisting of ST34Fs (‘CD34+Flk2- ST-HSC’) and STSLs (‘short-term reconstituting stem cell’), whose numbers were slightly reduced or increased upon FACT loss, respectively (Fig.4E, F). Lineage-committed progenitors changed in different directions. For example, the number of multipotent progenitors (MPP34F, CD34+Flk2+ multipotent progenitors) was increased, while the number of committed progenitors changed either up (MLP – multilymphocyte progenitors, CMP – common myeloid progenitors, CDP - common dendritic cell progenitors, GMP - granulocyte-monocyte progenitor) or down (MEP – megakaryocyte- erythroid progenitors, MDP – monocyte dendritic cell progenitors) (Fig. 4E, F). Thus, the stem cells most sensitive to FACT depletion in bone marrow were the earliest hematopoietic progenitors, LTHSCs. Changes in the number of other stem cells may be direct (in case their viability depends on FACT) or secondary (redistribution in response to loss of cells dependent on FACT).

To understand the differences in the neutrophils (increased in clusters 1–3, 5; decreased in clusters 6–8), we ran Gene Set Enrichment Analysis (GSEA, MSigDB) using gene markers specific for these clusters (Fig. 4G and Table EV2). We found that FACT loss resulted in more neutrophils expressing genes of the Interferon (IFN) gamma response and induced by other inflammatory cytokines (e.g., TNF and IL-6), while genes expressed in neutrophils that decreased in number were enriched in gene lists for proliferative response and elevated metabolic turnover (Fig. 4F and Table EV2). Common genes between the two sets of clusters were enriched in several neutrophil datasets (Table EV2).

We ran the same analysis for B cells by comparing cluster 17 (increased) with clusters 18 and 19 (decreased). The total number of B cells increased almost two-fold (6.5% vs. 13.1%, Fig.4C), suggesting that they might migrate from outside the bone marrow. Results of this comparison paralleled the neutrophils (Fig.4H and Table EV2). Upon FACT loss, there were more B cells with a transcriptional program similar to ‘Allograft rejection’ and ‘IFN gamma response,’ while in the control bone marrow there were more B cells expressing genes enriched for ‘MYC targets,’ ‘G2M checkpoint, and ’ E2F targets’. Thus, these data suggest that FACT depletion in neutrophils and B cells resulted in the inhibition of the expression of genes involved in proliferation and metabolism and the induction of genes induced by IFNs and other inflammatory cytokines.

Intestine UMAP plots showed that three neighboring clusters were completely lost (Fig.5A, clusters 1–3), while several clusters increased (clusters 10 - 14) or emerged de novo (clusters 4, 8, 9, 16, which had no or a very few cells in control sample). To recognize intestinal cells, we used data published by Gao et al. [29] and Haber et al. [30]. All clusters were ascribed to certain intestinal cell types, except cluster 12, which was not identified (Fig. 5B and Table EV3). In most cases, comparison of our data with both published datasets resulted in assignment of cells to the same type. The transcription signature of the most prominently lost cluster (1) included ‘large intestine Lgr5-positive stem cells’ [29]. Cells of this cluster also had high expression of *Gkn3*, a marker of small intestine stem cells [30] (Fig. 5C, D).

**Figure 5.**
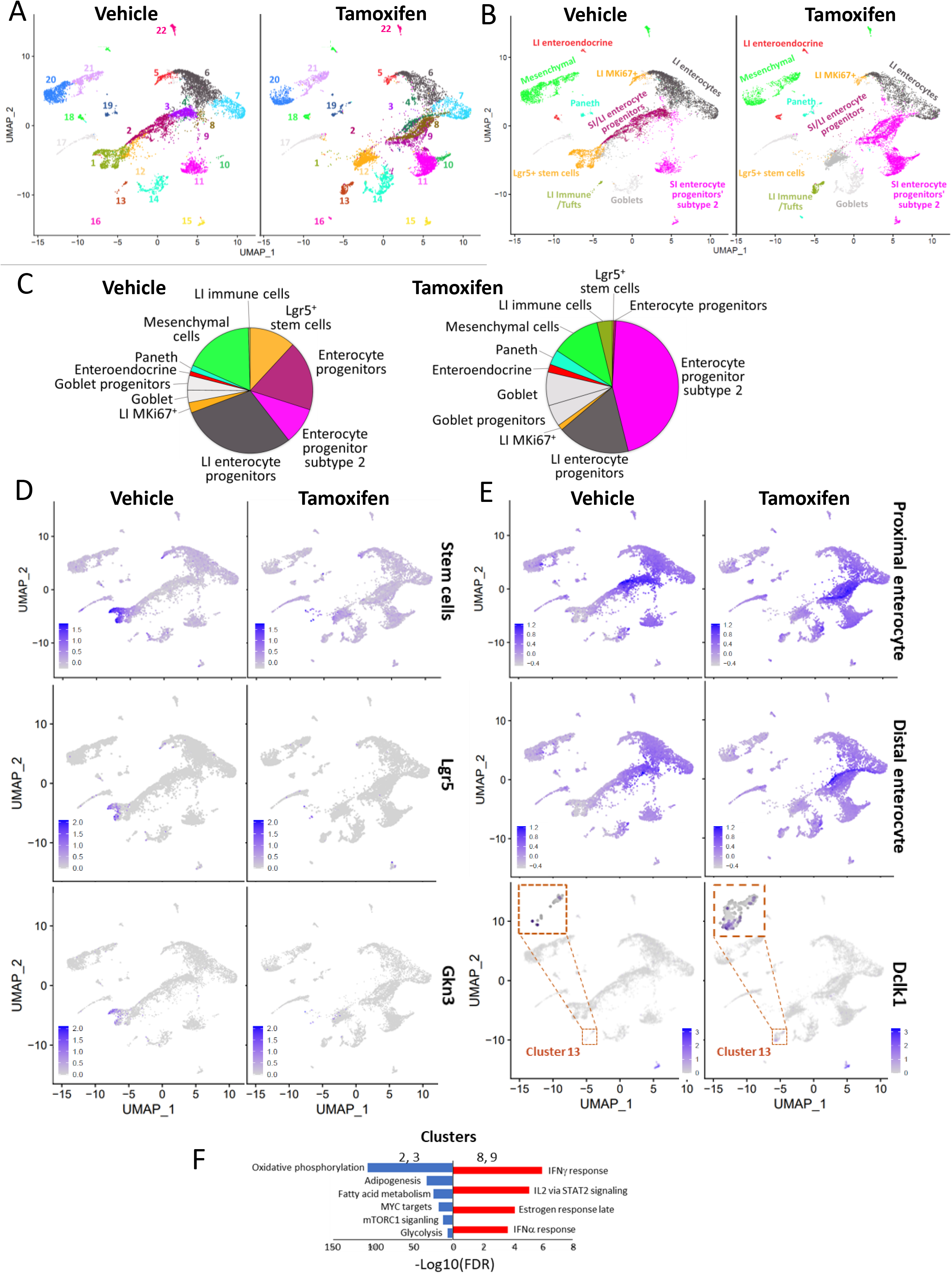
Effect of FACT depletion on cell composition of the intestine from *Ssrp1^fl/fl^; CreER^T2+/+^* mice (pooled data from two mice on day 1 after the end of treatment). A. UMAP plots with unbiasedly identified clusters shown in different colors. B. UMAP plots are colored according to cell type identified by transcriptional signatures using data from [29] as a reference. LI – large intestine, SI – small intestine. C. Distribution of intestinal cell types in samples from vehicle- and tamoxifen-treated mice. Pie charts are colored accordingly to panel B. D. UMAP plots with color-coded intestinal stem cells identified by transcriptional signatures of intestinal stem cells from [30] or by expression of *Lgr5* or *Gkn3* genes. E. UMAP plots showing the distribution of cells with transcriptional signatures of proximal and distal enterocytes from [30] and cells expressing the *Dclk1* gene. F. GSEA of gene markers of clusters 2 and 3 (lost upon *Ssrp1* KO, blue) versus clusters 8 and 9 (emerged upon *Ssrp1* KO, red).

No other cell types were completely lost upon FACT depletion. Lost clusters 2 and 3 were recognized as small and large intestine enterocyte progenitors using the Gao dataset (Fig. 5B, C) or proximal enterocytes using the Haber dataset (Fig. 5E). However, in the FACT-depleted intestine, similar cells were found within de novo emerged clusters 4, 8, and the increased clusters 9 –11 (Fig.5B, C, E). The close location of the reduced and increased clusters and their similar classification suggested that the enterocyte progenitors did not die upon FACT loss but had their transcriptional program shifted. Thus, we looked for differences in gene markers of clusters 2 and 3 (almost completely lost in the *Ssrp1* KO sample) and clusters 8 and 9 (emerged in the *Ssrp1* KO sample) (Fig. 5F) using GSEA. This comparison showed that FACT depletion in enterocyte progenitors resulted in downregulation of genes involved in ‘oxidative phosphorylation,’ ‘adipogenesis,’ ’ mTORC1 signaling,’ and ‘glycolysis,’ or being ‘MYC targets, while the upregulated genes were part of categories ‘IFN gamma and alpha response,’ ‘activated by IL2 via STAT2 signaling,’ and ‘late estrogen response’ (Fig. 5F and Table EV3). Surprisingly, these results were similar to what we observed with bone marrow neutrophils and B cells.

There were little changes in the number of cells in clusters 6 and 7 (Fig.5A), composed of distal enterocytes according to Haber or large intestinal enterocytes according to Gao. Quantitation revealed a slight increase in the number of Goblet cells and their progenitors and Paneth cells (Fig. 5B, C). Cluster 13 had an increased number of cells upon FACT loss (Fig.5A). These cells had a gene expression signature for large intestine immune cells (FDR q-value = 9.93 e-65) and were positive for Dclk1, a marker of intestinal Tuft cells (Fig.5E). Changes in intestinal cell composition were similar to those seen in bone marrow, namely the disappearance of the earliest intestinal stem cells and an increase in the number of more mature cell types. Cells in a state between stem and fully differentiated likely respond to FACT loss by reducing proliferation and inducing the IFN response.

Thus, scRNA-seq data suggest that the cell types most sensitive to FACT loss are the undifferentiated stem cells. More mature differentiated cells tolerate FACT depletion; however, these findings need to be confirmed by additional methods.

#### Stem cells from several organs cannot expand *in vitro* in the absence of FACT

To confirm that stem cells are dependent on FACT for viability and growth, we compared the colony-forming ability of stem cells from FACT-sensitive organs. Cells isolated from bone marrow, small intestine, and colon were plated in stem cell medium and treated with 4- hydroxytamoxifen (4-OHT, active metabolite of tamoxifen) to induce *Ssrp1* deletion *in vitro*. A similar experiment was performed with mesenchymal stem cells (MSCs) from adipose tissue to determine whether the loss of adipose tissue is a primary or secondary effect of *Ssrp1* KO. While the growth of skin fibroblasts from the tails of the same mice was not affected by *Ssrp1* deletion [12], the growth of all stem cells was significantly reduced. No colonies formed from bone marrow isolated from *Ssrp1^fl/fl^; CreER^T2^* mice in the presence of 4-OHT; however, 4-OHT did not interfere with the growth of bone marrow cells from genotypes not resulting in bi-allelic *Ssrp1* deletions (Fig.6A, B). No organoids formed from the small intestine or colon stem cells from *Ssrp1^fl/fl^; CreER^T2^* mice upon 4-OHT administration (Fig.6C, D). The adipose MSCs were growing in medium with 4-OHT but at a slower rate than the vehicle-treated cells, and their numbers were reduced due to increased apoptosis upon FACT depletion (Fig.6E, H).

**Figure 6.**
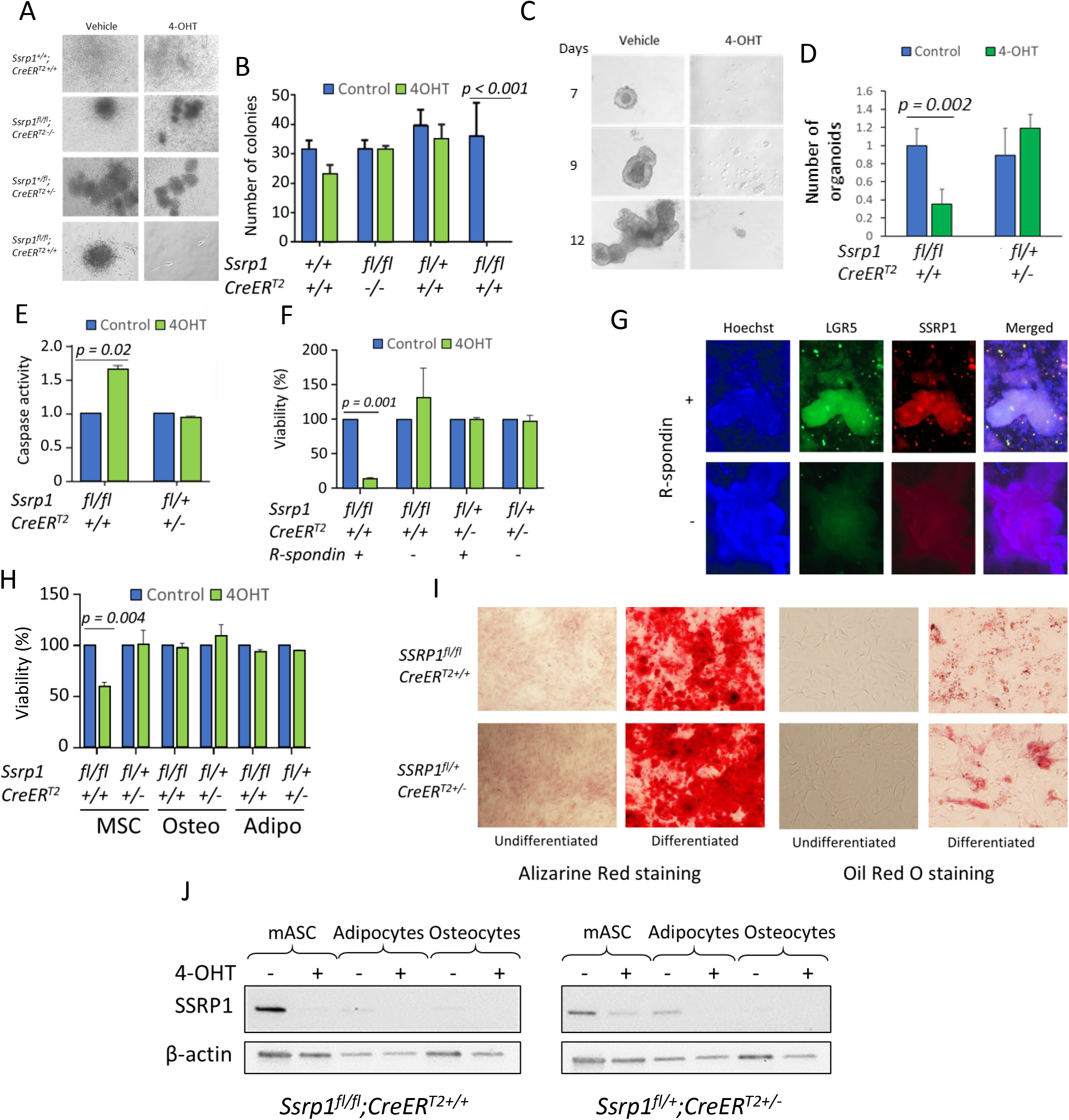
FACT depletion affects the viability of stem cells but not differentiated cells. A, B. Effect of 4-OHT administration on colony formation by bone marrow cells of different genotypes*. A.* Photographs of colonies from bone marrow cells kept in culture for seven days in the presence or absence of 4-OHT. B. Quantitation of bone marrow colony assay. Mean colony number per plate ± SD, *n = 3 plates*. P values < 0.05 are shown (unpaired t-test). 2 mice of each genotype were analyzed. C, D. Growth of intestinal organoids in the presence or absence of 4-OHT. C. Photographs of small intestine organoids on different days after plating. D. Number of organoids in the presence and absence of 4-OHT. Mean fold change between number of organoids in control and 4-OHT treated wells ± SD, *n = 4 wells*. P < 0.05 are shown (unpaired t-test). Organoids were isolated from one mouse of each genotype in two independent experiments. E. Caspase activity in lysates of control and 4-OHT treated MSCs of different genotypes. Mean fold change ± SD, *n = 3 plates*. P < 0.05 are shown (unpaired t-test). MSCs were isolated from 2-3 mice of each genotype. F, G. Effect of 4-OHT administration on intestinal organoids of different genotypes kept in the presence (undifferentiated) or absence (differentiated) of R-spondin-1. F. The viability of organoids assessed by the resorufin assay. Mean ± SD, *n = 3 wells*. P < 0.05 are shown (unpaired t-test). Organoids were isolated from two mice of each genotype. G. IF images of intestinal organoids from *Ssrp1^fl/fl^; CreER^T2+/+^* mice kept in the presence or absence of R-Spondin-1. The organoids were stained with antibodies to LGR5 and SSRP1 and counterstained with Hoechst (DNA). H–J. Effect of 4-OHT administration on MSCs and their differentiated counterparts. H. Methylene blue viability assay for undifferentiated MSCs or MSCs differentiated into osteocytes or adipocytes for 21 days and then treated with vehicle or 4-OHT. Mean ± SD, n = 3 plates. P < 0.05 are shown (unpaired t-test). MSCs were isolated from two mice of each genotype. H. Staining of MSCs, osteocytes, and adipocytes with Alizarine Red (osteocytes) or Oil Red (adipocytes). J. Western blotting of protein extracts from MSCs, adipocytes, and osteocytes from mice of different genotypes treated with vehicle or 4-OHT.

To determine if cells lose their dependence on FACT for viability upon differentiation, we isolated intestinal and adipose stem cells and grew them under conditions that either maintained their undifferentiated state or induced differentiation *in vitro* [31–33]. For both types of cells, 4-OHT administration was toxic only to undifferentiated cells. Intestinal organoids maintained in the absence of R-spondin 1 differentiated into more mature enterocyte-like cells [31], lost expression of *Lgr5* and became insensitive to FACT depletion (Fig.6 F, G). In the case of the adipose MSCs, we obtained cultures of differentiated adipocytes and osteocytes using specific differentiation media [34] and demonstrated that neither adipocytes nor osteocytes died upon FACT depletion (Fig.6H, I). Thus, only undifferentiated stem cells are dependent on FACT for viability. Consistent with this data, FACT levels decrease upon differentiation (Fig. 6G, J).

### 5. FACT loss in stem cells leads to significant changes in chromatin accessibility in a transcription-dependent manner

To understand the cause of stem cell loss upon FACT depletion, we investigated the changes in transcription and chromatin structure in adipose MSCs, cells which viability and growth depend on FACT, but which do not die upon FACT loss. First, we compared general transcription in MSCs before and after FACT depletion by measuring the incorporation of modified ribonucleotide 5-Ethynyl Uridine (EU) into newly synthesized RNA. We observed a slight but highly reproducible increase in EU incorporation in cells without FACT compared to those expressing FACT (Fig. 7A, B), suggesting that general transcription was increased in the MSCs upon FACT loss. This phenomenon was not the result of 4-OHT treatment since cells heterozygous for the *Ssrp1^fl^* allele showed almost identical EU incorporation before and after treatment (Fig. EV4A, B).

**Figure 7.**
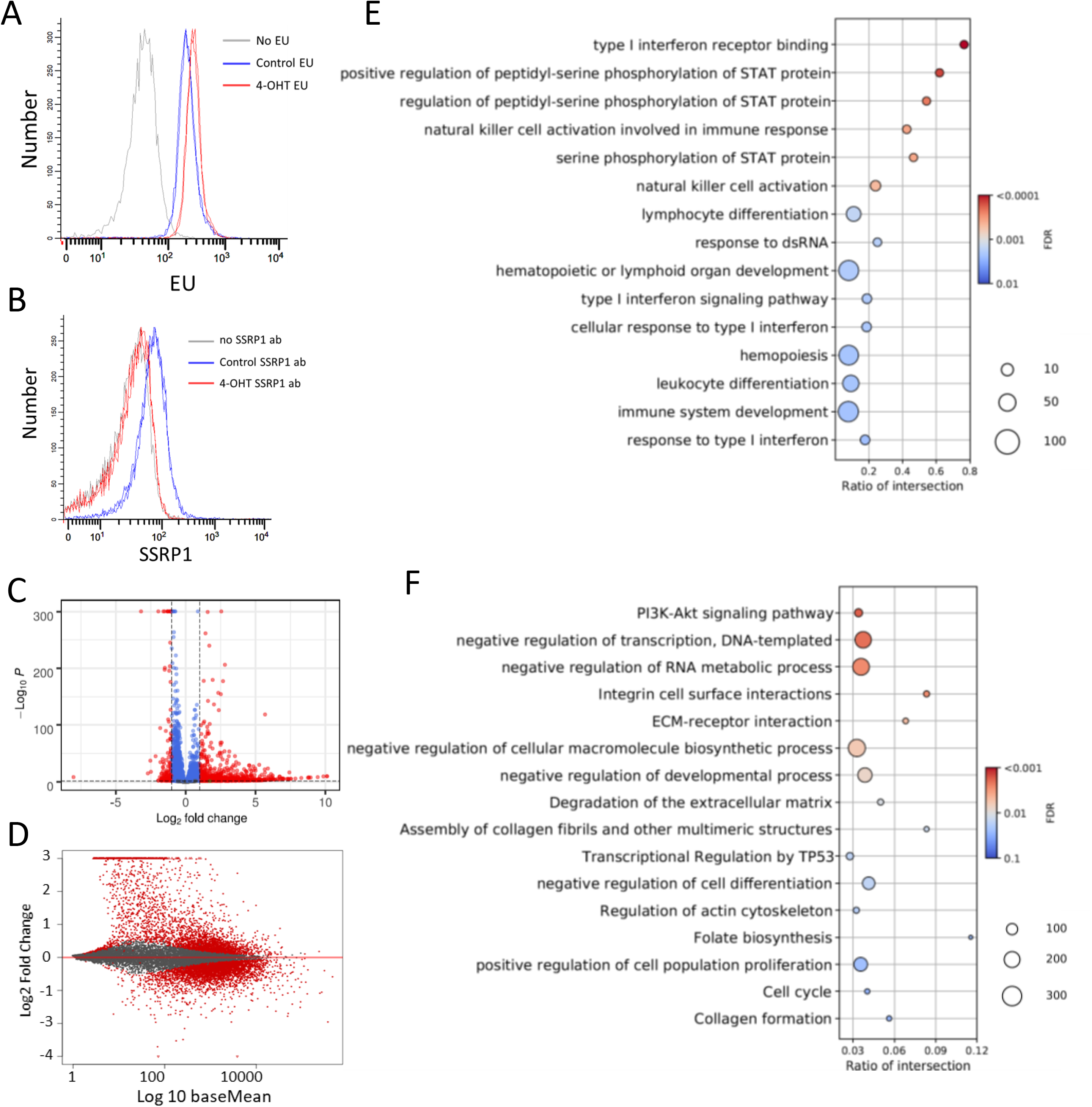
Comparison of transcription between *Ssrp1^fl/fl^; CreER^T2+/+^* MSCs treated with vehicle or 4-OHT. A. EU incorporation measured by flow cytometry. Histograms of EU fluorescent signals from two replicates for each condition (except no EU). B. SSRP1 staining of the cells in A. C. Volcano plot of changes in transcript counts in 4-OHT-treated cells versus control. Red dots indicate transcripts with fold-change ≥ 2 and FDR < 0.05. *n = 2 cell cultures*. B. MA plot showing the difference between the changes in transcript counts in 4-OHT-treated MSCs vs. control and the average expression of the same transcripts between conditions (baseMean). *n = 2 cell cultures*. C, D. Dot plots with the results of GO enrichment for genes upregulated (C) and downregulated (D) upon FACT depletion.

Next, we performed RNA-seq on samples from MSCs before and after 4-OHT treatment. There was a smaller number of genes for which expression changed in *Ssrp1^fl/+^* heterozygous cells than in homozygous *Ssrp1^fl/fl^* cells. There was almost no overlap in the genes whose expression was changed between the homozygous and heterozygous cells in response to 4-OHT (Fig. EV4C), confirming that the gene expression changes in the *Ssrp1^fl/fl^* cells were in response to FACT loss and not 4-OHT administration. In the homozygous cells, many more genes were upregulated upon FACT loss than downregulated (Fig.7C, D), whereas the changes in the gene expression of heterozygous cells were more random (Fig. EV4C, D).

Gene ontology (GO) analysis of the upregulated genes showed that the most significantly enriched category was the interferon response (Fig.7E and Table EV4). The downregulated genes did not have a dominant category but belonged to the ‘PI3K-AKT signaling pathway,’ ‘negative regulation of transcription,’ ’extracellular matrix (ECM) remodeling,’ ‘negative regulation of cell differentiation,’ and ‘positive regulation of cell proliferation’ categories (Fig.7F and Table EV4). Many of these categories reflected the observed phenotypic changes in these cells.

The chromatin state in the MSCs before and after 4-OHT treatment was analyzed using the Assay for Transposase-Accessible Chromatin with high-throughput sequencing (ATAC-seq) [35]. Because cells in different phases of the cell cycle have different degrees of chromatin accessibility, we limited the influence of this factor by maintaining cells for 48 hours in serum- depleted medium after 4-OHT treatment. Profile plots were built for the distribution of ATAC- seq reads at genomic regions flanking all known mouse genes, 3 kb before the transcriptional start site (TSS), 3 kb after the transcription end site (TES), and in between. Transposase inserts adaptors into genomic regions lacking nucleosomes (i.e., nucleosome-free or linker DNA of accessible chromatin). These adapters can be used as primers for PCR amplification. Selection of the length of the amplified fragments allows the identification of nucleosome-free regions (i.e., reads with the length shorter than nucleosomal DNA [< 120 bp]) or regions occupied by one nucleosome (i.e., reads with fragment lengths corresponding to mononucleosomes with linker DNA [>120 and <250 bp]) as described by Buenristro [35]. The profile plots showed very good overlap between replicates of the vehicle- and 4-OHT-treated cells (Fig.8), suggesting high reproducibility of the effect. The plots built from the nucleosome-free and mononucleosome fragments showed increased chromatin accessibility upon FACT depletion around the TSS and at gene bodies (Fig.8A, B). The increase in the gene bodies was more pronounced for the mononucleosome fragments than the nucleosome-free fragments (Fig. 8C, D).

**Figure 8.**
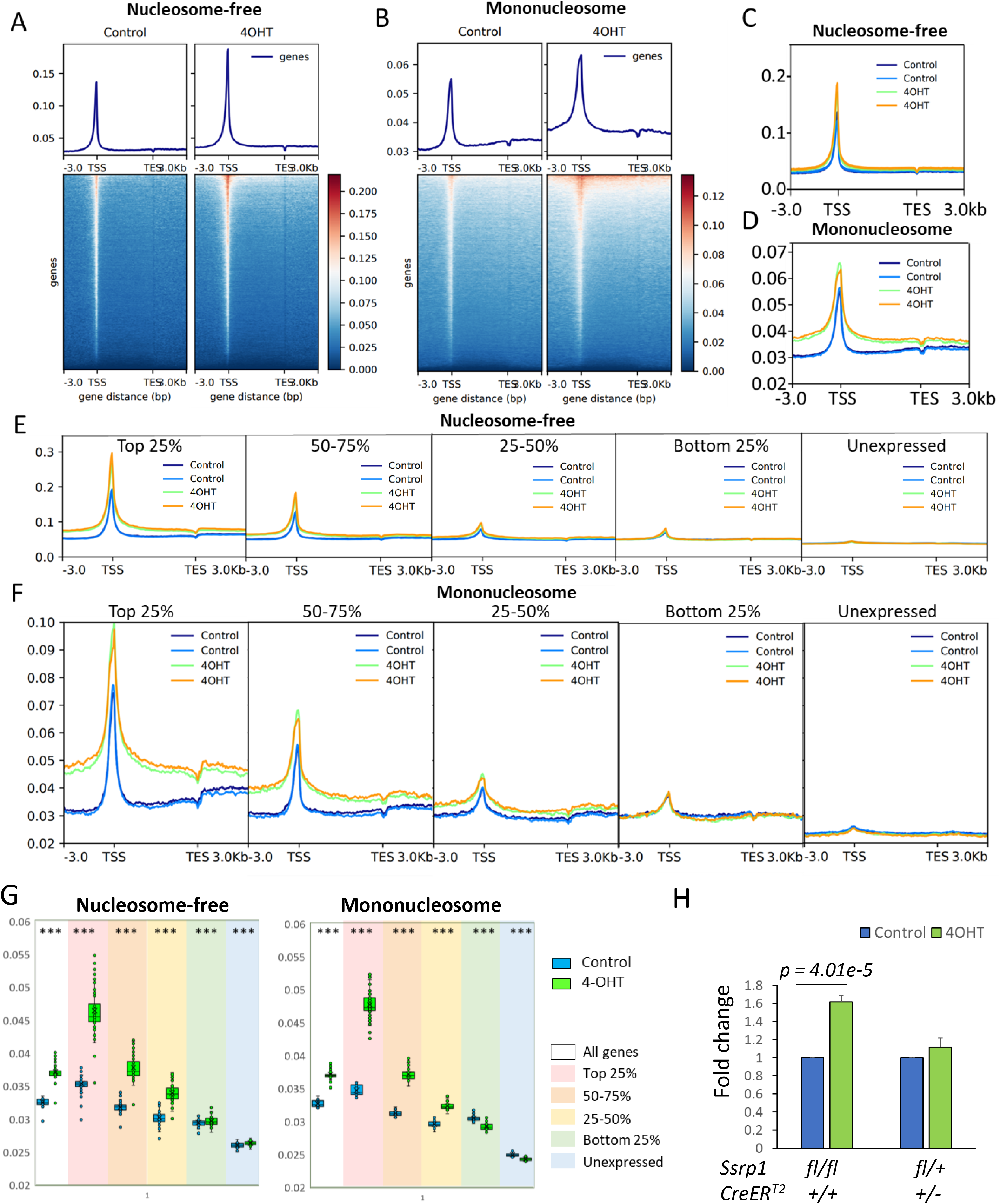
FACT depletion in *Ssrp1^fl/fl^; CreER^T2+/+^* MSCs leads to increase in chromatin accessibility in a transcription-dependent manner. A–B. Profiles and heatmaps for normalized ATAC-seq read distribution averaged for all known mouse genes in control or 4-OHT-treated MSCs for reads below 120 bp (A, nucleosome-free) and between 120 and 250 bp (B, mononucleosomes). One of two replicates is shown. C–D. ATAC-seq read density profile for two replicates of control and treated MSCs for all mouse genes of the nucleosome-free (C) and mononucleosome (D) fragments. E–F. ATAC-seq read density profiles for nucleosome-free (E) and mononucleosome (F) fragments for differentially expressed genes. G. Box plots of mean read density at the gene body (bins 21–100 out of 100 between TSS – TES) for all genes and differentially expressed genes. *** - p < 1e-10 (paired t- test). *n = 2 cell cultures*. H. Comparison of Hoechst 33342 binding to live control and 4-OHT-treated MSCs from different genotypes. Data are presented as the median fluorescence intensity ± SD, *n = 4 cell cultures*. P < 0.05 is shown (unpaired t-test).

To test whether FACT-mediated control of chromatin integrity depended on transcription, we used RNA-seq data to classify the genes based on their expression levels in cells with FACT. First, we identified genes that were not expressed in any of the tested conditions, which consisted of 26,577 genes out of the 55,401 total mouse genes analyzed. The remaining 28,824 genes were divided into four quantiles based on their average expression levels in cells with FACT. The most dramatic change was seen with the top 25% of the expressed genes, for which chromatin accessibility was significantly increased around the TSS and at gene body (Fig.7D). The differences were gradually reduced from the 5’ to 3’ ends and behind the TES. The differences were also reduced with the reduction in transcription but remained significant for the three top quantiles of the transcribed genes independent of whether nucleosome-free or mononucleosome fragments were analyzed (Fig.8E, F).

The differences in chromatin accessibility were not seen for the bottom 25% of the expressed and unexpressed genes when we analyzed the nucleosome-free fragments, while the plots for the mononucleosome fragments for the same categories of genes at the gene bodies were even reverted (i.e., FACT loss caused a slight reduction in chromatin accessibility at the gene bodies with very low or no transcription). To quantify these effects, we compared the mean read density at the gene bodies (excluding the region immediately after the TSS; see the Materials and Methods for details) for the different categories of genes and fragment lengths (Fig.8G). This analysis confirmed there were fewer mononucleosome fragments at poorly transcribed genes upon FACT loss.

Untranscribed genes constitute more than half of all the genes used for the analysis and may represent other non-transcribed regions that constitute most of the genome in mammals. Thus, we used a Hoechst 33342 to measure the overall effect of FACT loss on chromatin accessibility. Hoechst 33342 is a cell-permeable minor groove binding molecule whose fluorescence is many-fold increased when bound to DNA. An access of small molecules to the minor groove of DNA is limited when DNA is wrapped around nucleosomes [36, 37]. We incubated FACT-positive and negative adipose MSC cells with Hoechst 33342 and then measured the fluorescence of individual cells using flow cytometry. To avoid potential differences in the amount of DNA per cell due to different cell cycle phases, we arrested all cells in G1 by serum withdrawal for 48 hours. As a positive control, we used the small molecule curaxin CBL0137, which disrupts DNA/histone interactions in cells [9]. As expected, curaxin treatment significantly increased the fluorescence of both homozygous and heterozygous cells (Fig. EV5A). The same effect was observed after 4-OHT treatment of homozygous but not heterozygous cells (Fig.8H and EV5B).

Thus, FACT loss resulted in increased chromatin accessibility, the degree of which was dependent on the level of transcription. The increase in both nucleosome-free and mononucleosomal fragments was likely due to the opening or partial unwrapping and loss of nucleosomes.

## Discussion

Transcription research has mainly focused on how RNA polymerase accesses nucleosomal DNA and what factors facilitate this process. FACT has been viewed as a factor facilitating transcription through chromatin [38]; however, accumulating data questions FACT’s role in opening nucleosomes to RNA polymerase (reviewed in [39, 40]). These data may be summarized as follows: (i) there is no evidence that FACT removal from eukaryotic cells results in inhibition of transcription, while it has been shown that FACT depletion in several model systems results in no change or even an increase in the absolute transcription levels [12, 16]; (ii) there is no strong data that FACT can bind and destabilize nucleosomes with fully wrapped DNA; however, there is data indicating that FACT binds all nucleosomal components and nucleosomes not fully wrapped with DNA (reviewed in [40]); (iii) decreased FACT levels in cells results in loss of histones from chromatin [41–43] and increased accessibility of chromatin [11]. These observations led to the emergence of a new model for the role of FACT in transcription: FACT binds the components of nucleosomes, histone dimers/tetramers, and possibly DNA, to preserve nucleosomes at their positions during RNA polymerase passage; without FACT, nucleosomes are prone to disassembly by transcription machinery or associated factors. This model can also explain the initially puzzling observation that while FACT enrichment at mammalian genes is highly proportional to the transcription level, the transcription of genes bound by FACT is not inhibited by FACT removal [12, 15, 44]. The greater the transcription levels (i.e., more frequent passage of RNA polymerase through a region), the more frequently nucleosomes are disturbed and the more opportunities for FACT to bind. As previously demonstrated, the re-establishment of histone-DNA contacts displaces FACT because of the higher affinity of histones for DNA than for FACT [45]. Therefore, genes with lower transcription (i.e., less frequent passage of RNA polymerase) accumulate less FACT. At the same time, if FACT loss is accompanied by the loss of nucleosomes from the transcribed region, we might expect an facilitated passage of RNA polymerase through this region in the absence of nucleosomes, i.e. elevated transcription.

In this study, we tested and confirmed this proposed model in a mammalian system. Although in this study we genetically excised only *Ssrp1* gene, we have never observed in mammalian cells existence of SPT16 in the absence of SSRP1 and unbound to SSRP1. Thus, we discuss below the consequences of the loss of the whole complex rather than SSRP1 subunit, though theoretically SSRP1 may have independent activity in some cells or conditions.

We demonstrated that FACT is the major factor responsible for the maintenance of chromatin during transcription and is essential for the viability of a small fraction of cells in adult mice. However, these cells are vital to the viability of adult mice because *Ssrp1* KO was lethal for mice at all tested ages (5–40 weeks). Together with previously published data that general *Ssrp1* KO is lethal at 3.5 dpc [18], our new data revealed that *Ssrp1* is an essential gene in mammals because it is critical for the survival of stem cells. Since we did not use aged mice in our experiments, consequences of FACT loss in geriatric animals requires further investigation.

The hematopoietic and intestinal tissues were the most sensitive to FACT loss. Both tissue types are dependent on the constant replenishing of cells from tissue-specific stem cells. The earliest progenitors in both systems were completely lost upon *Ssrp1* KO. The stem cells in other tissues may also be sensitive to FACT loss, as indicated by the depletion of adipose tissue and the presence of skin abnormalities. For testing the role of FACT in the viability of other tissues, tissue-specific *Ssrp1* KO is needed to prevent the death of animals from the bone marrow and intestinal failure.

FACT is not essential for many types of adult cells [12, 16]. Here, we established that number of many cell types, mostly differentiated but also some progenitors, increased upon FACT loss. This increase is likely due to FACT-independent, compensatory redistribution of cell populations; however, time-dependent monitoring of organ cell composition upon *Ssrp1* KO is needed to establish this. In addition, the abundance of these cell types, even among stem and progenitor cells, suggests that multiple adult cells do not need FACT for survival. Importantly, there were many proliferating cells among the FACT-independent cell types. We also did not see changes in the cell-cycle distribution upon FACT loss, suggesting that stem cells disappear not due to problems with replication or mitosis, processes where FACT activity has been observed [23, 27]. Thus, an important question is what makes these cells sensitive to FACT loss. To address this question, we showed that FACT loss resulted in a significant increase in chromatin accessibility. Previously, we observed that FACT removal from differentiated fibroblasts, which express low FACT levels and are not sensitive to *Ssrp1* KO, does not lead to the loss of nucleosomes, while it does in oncogene-transformed cells [12]. Therefore, we propose that there is a specific chromatin state in some stem cells and oncogene-transformed cells, where nucleosomes are not as stable as in other cells what makes these nucleosomes as well as viability of cells dependent FACT.

There is a clear difference between the sensitivity of chromatin to FACT loss depending on transcription, i.e., the higher the transcription rate, the greater the increase in chromatin accessibility upon FACT removal for most of transcribed genes (top 75% of genes). These changes were in line with the increase in global transcription in MSCs upon FACT depletion. However, there was a small but highly significant reduction in chromatin accessibility at untranscribed genes or those with low transcription rates upon FACT loss (at least if comparing mononucleosome fragments). This finding could potentially be explained by the role of FACT in preventing the stalking of folded nucleosomes through binding to the acidic patches [46]. In this case, the loss of FACT in the absence of transcription might lead to a tighter interaction between nucleosomes, leaving fewer opportunities for transposase to invade the linker DNA.

An interesting observation that may shed light on the physiological consequences of chromatin destabilization is the increase in the number of immune cells in the bone marrow and intestine. The increase in B cells in bone marrow may be due to a shift in differentiation, whereas the number of T cells in the bone marrow and intestine may only be increased due to migration. This accumulation may be related to the fact that the most upregulated pathway in response to *Ssrp1* KO in all cell systems examined was the IFN response. We previously observed activation of the IFN response in mice treated with curaxin CBL0137, which disrupts histone/DNA binding [9]. IFNs are well-established factors that attract immune cells to the sites of viral invasion and inflammation [47]. The mechanism of interferon activation upon chromatin destabilization is not fully established; however, in the case of curaxin, it resulted from dsRNA accumulating in cells due to the transcription of pericentromeric repeats [48]. The same mechanism may be involved in the effects of FACT depletion along with other mechanisms, such as the emergence of cryptic initiation from uncapped RNA sensed by the RIG/MAVS5 pathway [49] or the binding of nuclear cGAS to DNA losing nucleosomes [50].

In this study, we established that FACT performs a unique function in the earliest progenitor stem cells in several organs of adult mammals. We propose that in these cells, FACT is critical for the maintenance of chromatin during transcription. Loss of FACT results in the loss of nucleosomes at transcribed genes and elevated transcription. The most significantly upregulated pathway in response to the increased chromatin accessibility is interferon signaling, which is likely responsible for the accumulation of immune cells in the sensitive organs in response to FACT loss.

## Materials and Methods

### Reagents

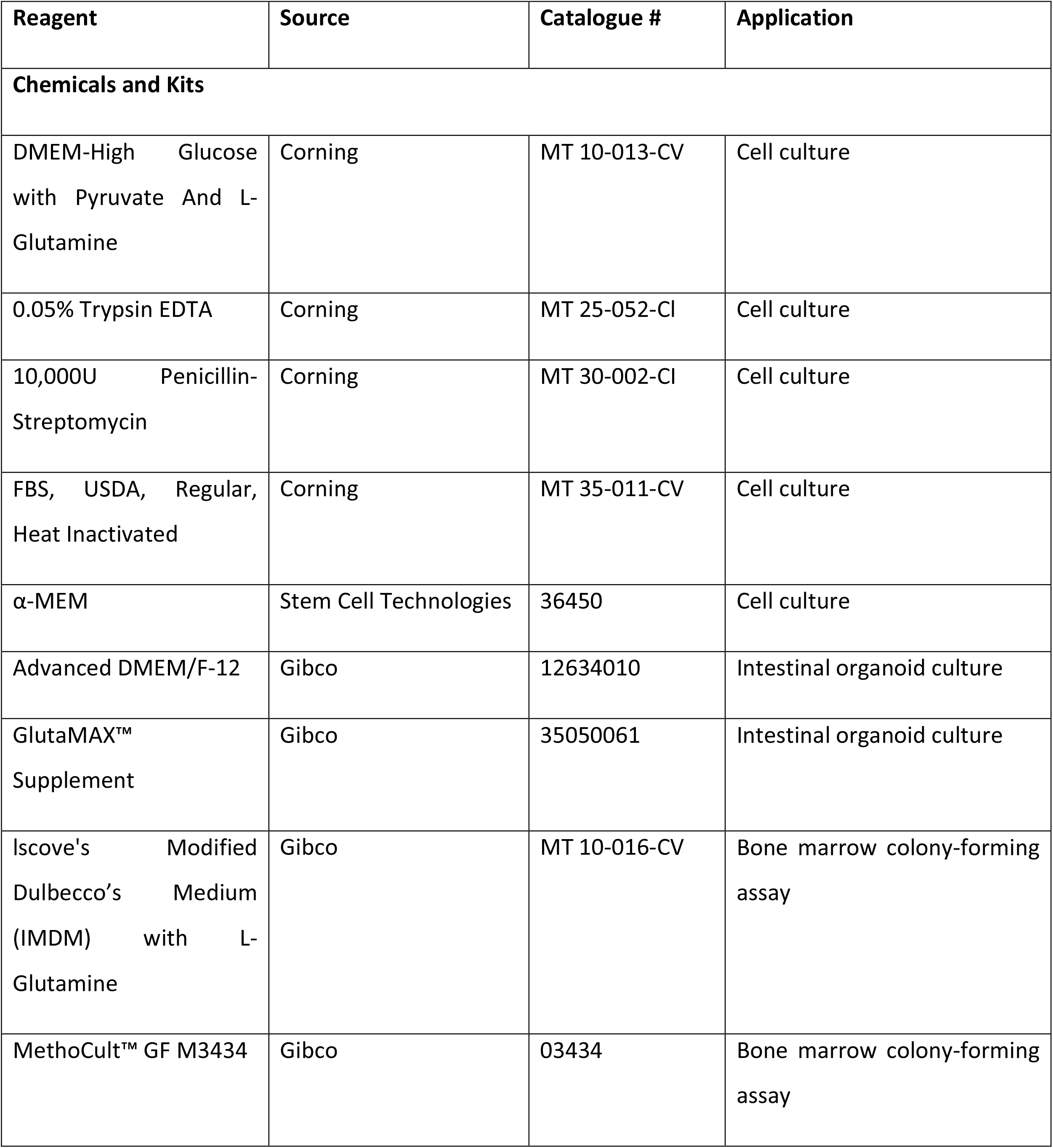
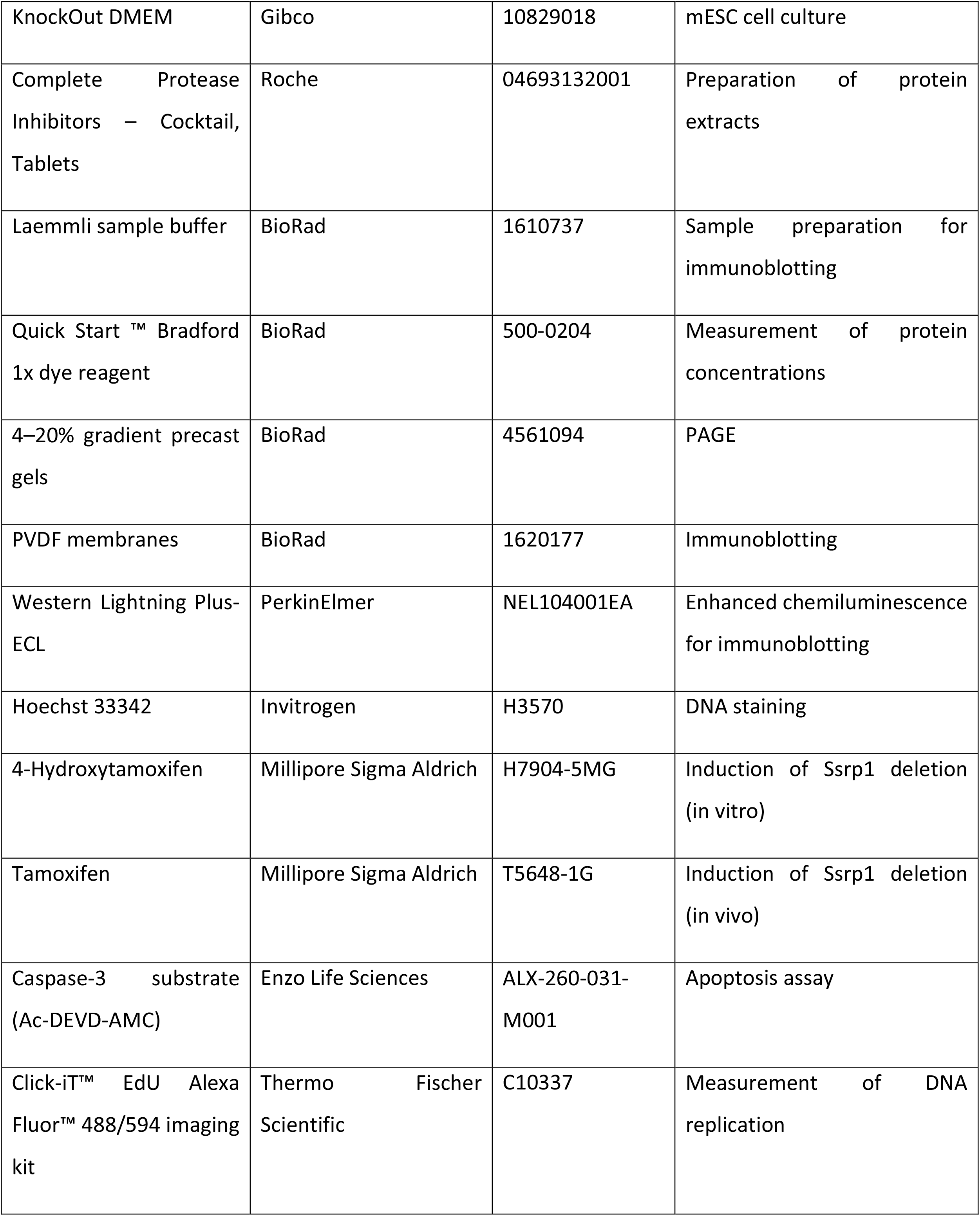
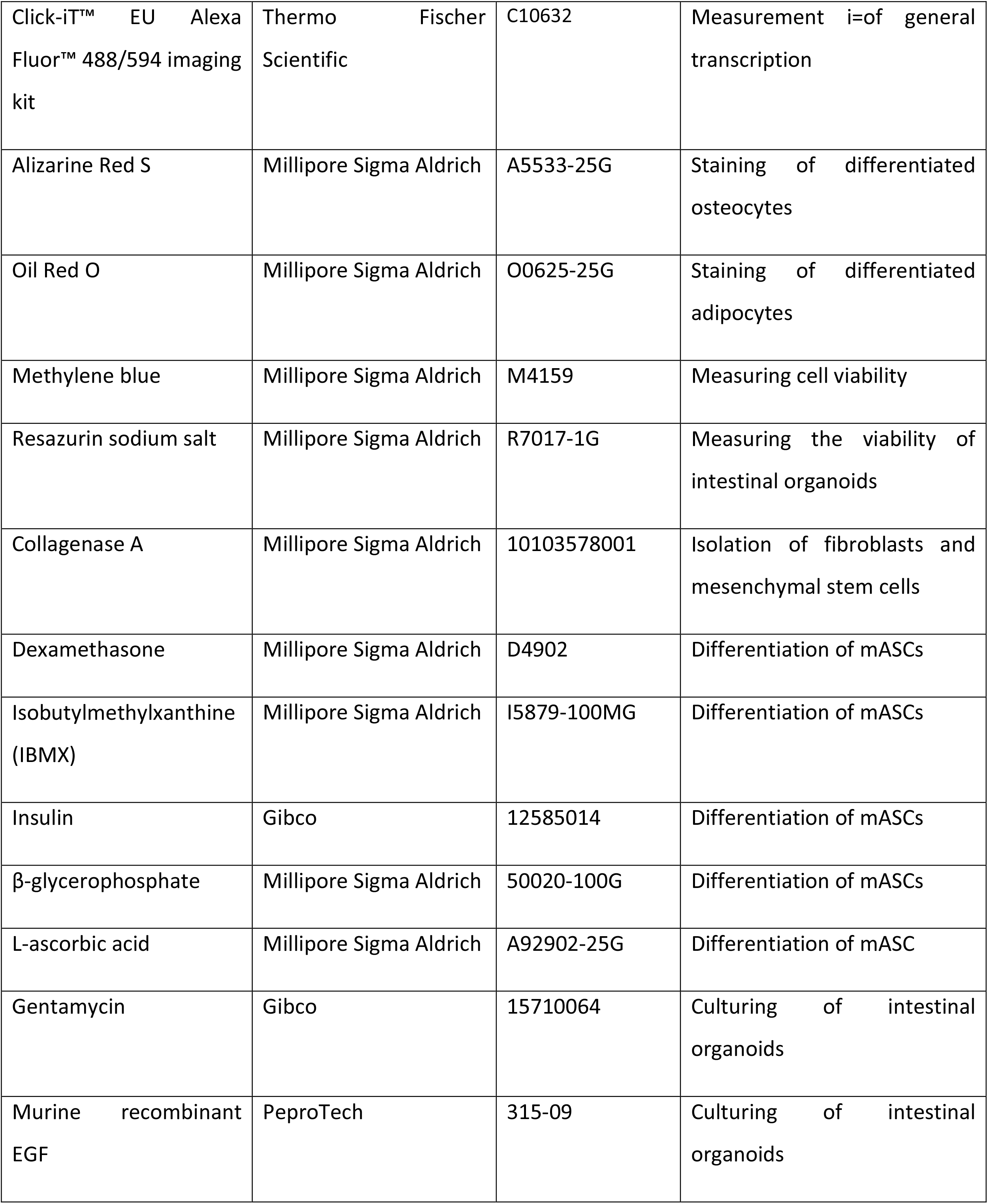
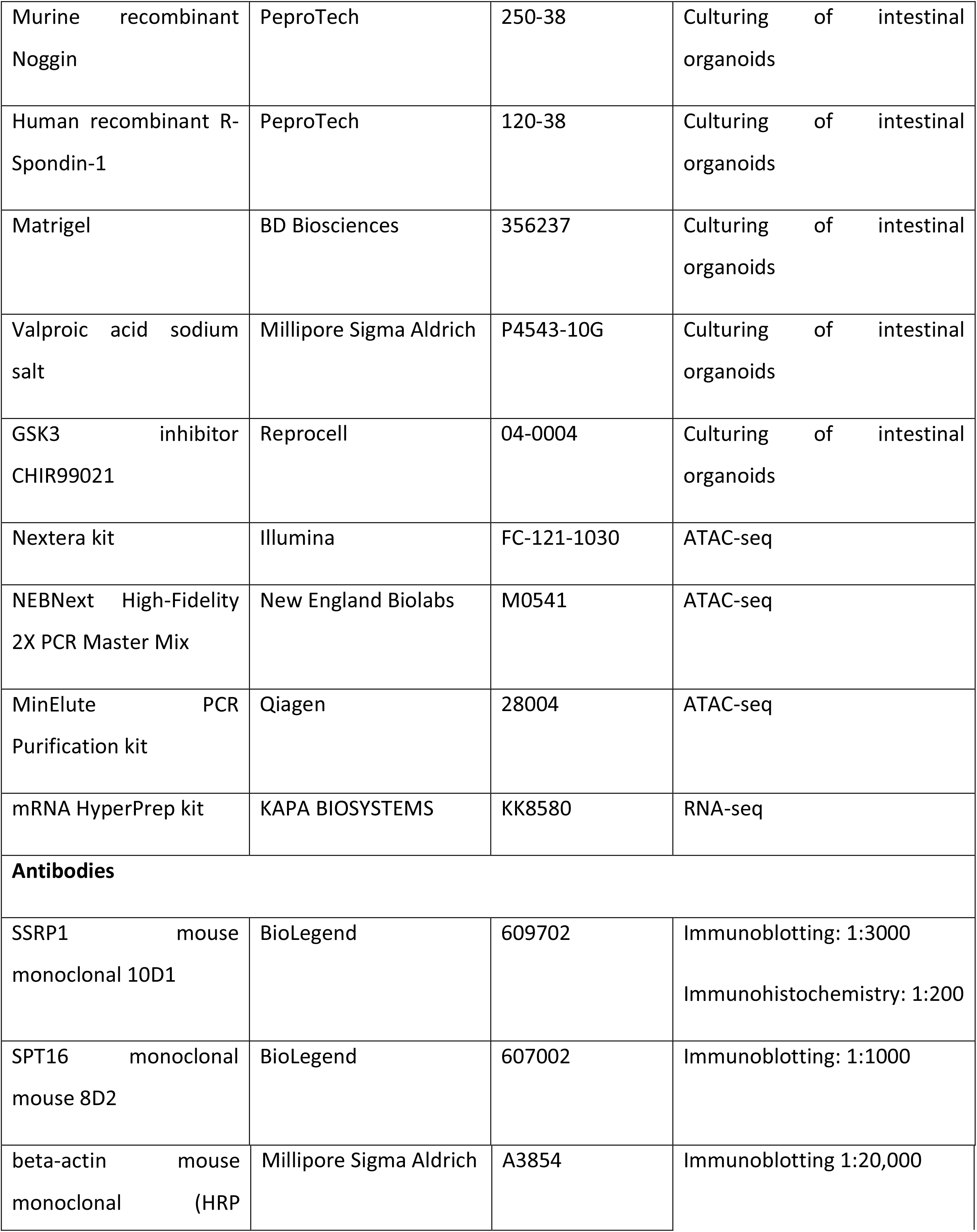
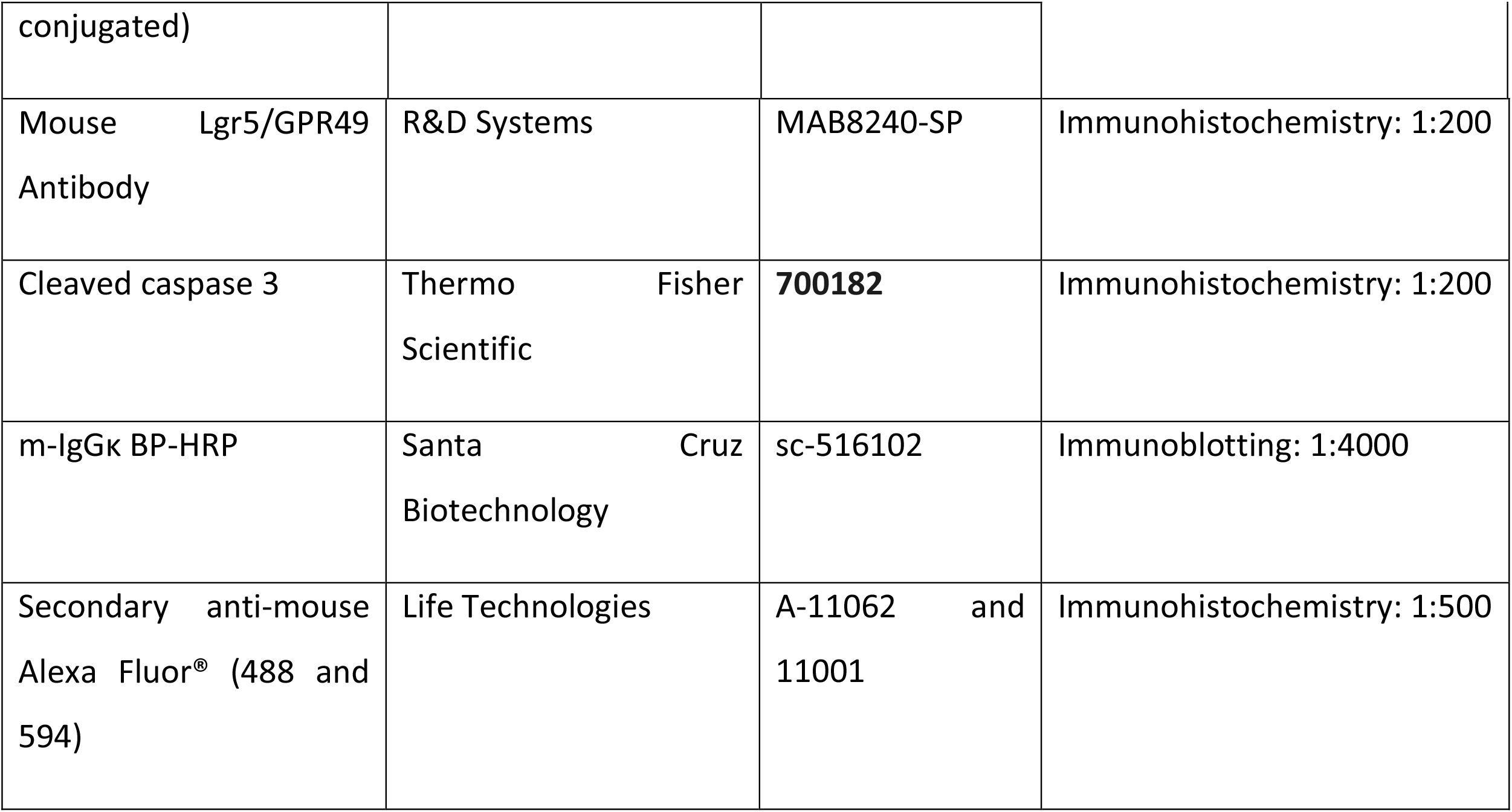

The 10 mg/mL tamoxifen solution was prepared by mixing 100 mg tamoxifen powder in 100 μL 99% proof ethanol and then slowly dissolving this in 9.9 mL sterile corn oil at 37⁰C overnight. A 5 mM stock solution of 4-hydroxytamoxifen (4-OHT) was prepared by dissolving 4.8 mg 4-OHT powder in 2.5 mL 99% proof ethanol.

Alizarin Red S was prepared by dissolving 2 g powder in 100 mL distilled water. The pH of the solution was adjusted to 4.1–4.3 with 10% ammonium hydroxide. The solution was stored at 4⁰C away from light.

Oil red O stock solution was prepared by dissolving 0.7 g powder in 200 mL isopropanol, stirring overnight, and filtering through a 0.2-μm filter. The solution was stored at 4⁰C away from light. The working solution was prepared fresh by diluting six parts oil red O stock into four parts distilled water. The mixture was incubated at room temperature for 20 min and filtered through a 0.2-μm filter.

A 2.5 mM resazurin solution (36X) was prepared by dissolving 0.629 g resazurin sodium salt (Milipore Sigma) in Dulbecco’s phosphate-buffered saline (DPBS) and sterilizing through a 0.22 μm filter. The solution was stored at -20⁰C protected from light. The working solution was prepared by diluting this stock to 6X. The working solution was stored at 4⁰C protected from light for up to 3 months. The 6X solution was used at a final concentration of 70 μM.

### Animal experiments

#### Animal breeding and treatment

All animal breeding, procedures, and experiments were conducted in accordance with the Institute Animal Care and Use Committee (IACUC) at Roswell Park Comprehensive Cancer Center under an approved IACUC protocol. Mice were maintained in the Laboratory Animal Shared Resource with controlled air, light, and temperature conditions and fed ad libitum with 5% fat chow with free access to water.

The conditional FACT Knockout mouse model with the following genotypes were generated as previously described [20]: *SSRP1^fl/fl;^ CreERT2^+/+^* (KO mice), SSRP*1^+/+^; CreERT2^-/-^* (wild type), *SSRP1^+/+^; CreERT2^+/+^*, *SSRP1^fl/fl^; CreERT2^-/-^*, *SSRP1^fl/fl^; CreERT2^-/-^*, and *SSRP1^fl/+^; CreERT2^+/-^* (controls).

Mice of different ages were treated with 100 μl of tamoxifen (1 mg) or vehicle by daily IP injection for five days. In some experiments, mice received daily subcutaneous injections of 0.9% saline (1 mL). Mouse weights and cage-side observations were recorded daily. Animals were euthanized by CO_2_ asphyxiation and cervical dislocation upon losing 20% of their body weight. All cells, tissues, and organs were harvested under sterile conditions.

#### Mouse tissue collection and staining

Mouse tissues were harvested and preserved in 10% buffered formalin. Paraffin-embedded tissue sections (5 μm) were stained with hematoxylin and eosin (H&E) and anti-SSRP1 and Anti-cleaved caspase-3 antibodies in the Immunohistochemistry Laboratory of the Roswell Park Pathology Resource Network using standard techniques. All slides were scanned using the Aperio system (Aperio Technologies, Inc). Images were collected using Image Scope software (Aperio Technologies, Inc).

EdU staining was performed three days after the end of tamoxifen treatment. Briefly, mice were injected with 50 mg/kg EdU (Invitrogen, Thermo Fisher Scientific) one hour before euthanasia. Tissue sections were stained using the Click-iT EdU Alexa 488 Fluor Imaging kit (Thermo Fisher Scientific) and counterstained with 4′,6-diamidino-2-phenylindole (DAPI) (Life Technologies) following the manufacturer’s instructions. The stained samples were examined using a Zeiss AxioImager A1 microscope equipped with an epifluorescent light source. Images were captured with an AxioCam MRc digital camera and processed with Zeiss Axio Imager Z1 microscope software (Carl Zeiss, Germany).

#### Magnetic resonance imaging (MRI)

Two mice of each gender and genotype were imaged. MRI was performed on day 0 before treatment and days 6, 8, and 10 after the start of treatment with tamoxifen or vehicle. Images were acquired on a 4.7 Tesla preclinical MRI scanner using the ParaVision 3.0.2 platform (Bruker Biospin, Billerica MA). A 72-mm (ID) quadrature RF coil was used to capture the volumetric data for the entire mouse body. Two NMR tubes filled with 1 mM and 3 mM CuSO4 in 1% agarose were included in each session for signal normalization. Following the scout scans, a three-dimensional, fast spin echo scan was acquired using the following parameters: TE/TR = 6.2/100 ms; echo train length = 4; averages = 4; FOV = 113 x 55 x 35 mm; matrix = 256 x 128 x128; zero-filled to achieve an isotropic voxel size of 273 microns. Image signal intensities were normalized by a three-point correction curve using two phantoms and background noise, and the whole-body volume was segmented and calculated using the region-growing tool in Analyze 7.0 software (AnalyzeDirect, Overland Park KS). Adipose tissue was segmented from lean tissue using a fixed intensity threshold derived from the average threshold of all datasets, as reported by the Otsu method (REF) and confirmed by visual inspection. The percent adipose tissue volume was calculated as a ratio of voxels segmented as adipose tissue divided by the total body voxels.

#### Complete blood counts (CBC) and serum biochemistry

Blood (50 μL) was collected from the saphenous veins of mice before treatment (baseline) and on days 7 and 12 after the start of tamoxifen treatment, as previously described [51]. For serum biochemistry, blood was collected from deeply anesthetized mice via cardiac puncture into serum separator tubes (Sarstedt). The samples were clotted at room temperature for at least 15 min and then centrifuged at 10,000 RCF for 10 min. The serum was analyzed using an IDEXX Catalyst DX Chemistry Analyzer (IDEXX Laboratories).

### Cells

#### Mesenchymal stem cells (MSCs)

MSCs were harvested from the adipose tissue of 8–10-week- old mice. The tissue was minced and digested for 30 min via shaking in a solution of 2 mg/mL collagenase A in PBS at 37⁰C. The digested tissue was filtered through a 40-μm strainer and centrifuged at 300 g for 5 min. The cells were resuspended in DMEM with 10% FBS and 1% penicillin-streptomycin and plated. To differentiate the MSCs into adipocytes and osteocytes, MSCs were plated on 60-mm plates. When cells reached 80% confluence, differentiation into adipocytes was induced by incubating the cells in medium containing 250 nM dexamethasone, 66 nM insulin, and 0.5 mM isobutylmethylxanthine (IBMX) for 21 days. Differentiation into osteocytes was induced by incubating in medium containing 0.1 μM dexamethasone, 10 mM β-glycerophosphate, and 50 uM L-ascorbic acid for 14 days. The differentiation media were changed every 2–3 days. Differentiation into adipocytes and osteocytes was determined by staining with Oil Red O and Alizarin Red, respectively. After differentiation, the cells were treated with 2 μM 4-OHT for five days, and viability was assessed by resorufin staining.

Staining for adipocyte and osteocyte differentiation (Oil Red O and Alizarin Red S, respectively) was done by removing the medium, washing cells with PBS, and fixing with 10% neutral buffered formalin for 30 min with gentle shaking at room temperature. The cells were washed twice with distilled water, and then 60% isopropanol was added to the cells for 5 min. The cells were incubated with the appropriate stain for 20–45 minutes at room temperature, protected from light. The plates were then washed twice with distilled water. The cells were covered with water, and images were taken with a light microscope.

#### Intestinal stem cell isolation and organoid culture

Intestinal stem cells were collected as previously described [52]. Briefly, approximately 15 mm of the intestines (small, immediately before cecum; large, immediately after cecum; cecum excluded) was harvested from mice and placed in cold PBS. The intestine was cut open longitudinally and flushed with cold PBS multiple times to remove all the contents. The tissue was cut into 5-mm pieces and incubated at 4⁰C in PBS containing 2 mM EDTA and 50 μg/mL gentamycin with gentle shaking for 30 min. The tissue was then resuspended in cold PBS and vigorously shaken to dislodge the crypts. After allowing the suspension to settle by gravity for 5 min, the supernatant containing the villus fraction was discarded. The tissue was resuspended in cold PBS and shaken vigorously to enrich the supernatant with the crypt fraction. The crypt fraction was filtered through a 70-μm strainer and centrifuged at 200 g for 3 min at 4⁰C. The pellet was resuspended in crypt culture medium (advanced DMEM:F12 containing 2% FBS, 50 μg/mL gentamycin, 2 mM L-glutamine, 50 ng/mL EGF, 500 ng/mL R-Spondin-1, 100 ng/mL Noggin, 1 mM valproic acid sodium salt, and 3 μM GSK3 inhibitor CHIR99021. The crypts were counted using a hemocytometer, and 500 crypts were mixed with 50 μL Matrigel at 4⁰C and pipetted into the center of the wells of 24-well plates prewarmed to 37⁰C. The domes were allowed to set for 10 min in a 37⁰C incubator, and then crypt culture medium (500 μL) was gently added to each well. Organoids formed over 7–10 days, with medium changed every 2–3 days. The crypts were passaged as previously described [52]. After two passages, the organoids were treated with 2 μM 4-OHT for five days. On day 6, the organoids were split into new wells and allowed to grow for 5–7 days. The ability to form colonies was assessed by counting the number of organoids in the control- and 4-OHT-treated wells. The data were normalized to the number of colonies formed in the control wells.

To differentiate the organoids, the intestinal stem cells were cultured in medium without R-Spondin-1 for 2.5 days as previously described [31]. The undifferentiated organoids were maintained in medium containing R-Spondin-1. The differentiated and undifferentiated organoids were treated with 2 μM 4-OHT for five days, and viability was tested at the end of treatment using resorufin staining.

For immunofluorescence, the organoids were fixed in 4% paraformaldehyde (PFA) for 30 min, which partly dissolved the Matrigel. The organoids were collected by centrifugation at 2000 rpm for 5 min at room temperature. The pellets were washed with PBS, and the organoids were permeabilized with PBS containing 0.1% Triton X-100 and 3% bovine serum albumin at room temperature with rotating. The samples were then blocked in PBS containing 0.05% Triton X-100 and 1% BSA for one hour with rotating. The organoids were stained with primary antibodies (SSRP1 [1:200], Lgr5 [1:200]) at 4⁰C overnight with rotating. The organoids were washed with blocking solution and incubated for 1 hr at room temperature with the following secondary antibodies: anti-rat Alexa Fluor 488 (1:500) against Lgr5 and anti-mouse Alexa Fluor 594 (1:500) against SSRP1. The organoids were washed with blocking solution and PBS and then counterstained with Hoechst (1:2000 in PBS) for 30 min at room temperature with rotation, protected from light. After staining, the organoids were washed with PBS and then plated in 96- well plates. Images were taken using a Zeiss Axio Imager Z1 microscope (Carl Zeiss, Germany).

#### Bone marrow colony-forming assay

The femurs and tibias of mice were excised from the hindlimbs by removing the muscles and connective tissue and cleaned with 70% ethanol. The bones were crushed using a sterile mortar and pestle in PBS to generate a single cell suspension that was filtered through a 70-μm cell strainer. This process was repeated until the bones appeared white, indicating all the marrow was removed. The cells were counted using a hemocytometer, and a suspension containing 4.5 x 10^5^ nucleated cells/mL was prepared.

Complete IMDM medium was prepared by adding 20 mL IMDM to 80 mL MethoCult M3231. 4- OHT (2 μM) or DMSO (control) was added to 4.1 mL complete IMDM by inverting the tube and vortexing for 5 sec twice. The tube was placed in a refrigerator for 5 min to allow bubbles to escape. The 4.5 x 10^5^ cells/mL bone marrow cell suspension (400 μL) was added to each tube of medium and the 4-OHT and control mixtures. The tubes were mixed gently by inverting the tube two times and vortexing three times for 8 sec each at maximum speed, followed by bubble release. The cell + medium + 4-OHT or control mixture (1 mL) was plated in 35-mm dishes using a primed 5-mL syringe with a 19G needle. The final cell density was 40,000 per plate. The plates were incubated at 37⁰C, and colonies were allowed to form on methylcellulose in the presence or absence of 4-OHT for seven days.

### Viability assays

For the methylene blue assay, 0.5% methylene blue in 50% methanol was added to plates after removing the medium. The plates were incubated for 15 min at room temperature with gentle shaking. The plates were washed with distilled water and air-dried. The dye was eluted for 30 min at room temperature with 1% sodium dodecyl sulfate (SDS) in PBS with shaking. The absorbance was read at 595 nm.

The resorufin assay was used to measure the viability of the intestinal organoids. Undifferentiated and differentiated Intestinal organoids were treated with or without 2 μM 4- OHT for five days in 96 well plates (100 μL). After treatment, 20 μL 6X resorufin solution was added to each well for a final concentration of 70 μM, and the plates were incubated at 37⁰C for 6 hr. The absorbance was read at 570 nm.

### EdU staining

Three days after the last treatment with vehicle or tamoxifen, 50 mg/kg EdU was administered intraperitoneally to the mice. After two hours, the mice were euthanized, and the tissues were harvested and snap-frozen. Tissue sections were fixed with 3.7% paraformaldehyde for 15 min at room temperature. For EdU treatment of cells, the cells were treated with 2 μM 4-OHT for five days and then split. After 24 hr, the cells were incubated with 20 μM EdU for 1 hr. The cells were then fixed with 3.7% PFA in PBS for 15 min at room temperature. Cells and tissue sections were permeabilized with 0.1% Triton X-100 in PBS for 20 min at room temperature. The incorporated EdU was detected using the Click-iT® Plus EdU Imaging Kit (Invitrogen), according to the manufacturer’s instructions.

### EU staining

MSC cells were treated with vehicle or tamoxifen for five days and then split. After 24 hr, the cells were treated with 0.2 mM EU for 1 hr. After the incubation, the cells were trypsinized, harvested, and washed with PBS in 1% BSA. The cells were fixed with 100 μL Click-iT fixative for 15 min at room temperature. The cells were stained with an SSRP1 antibody using a standard protocol and then for EU according to the manufacturer’s instructions. The cells were counterstained with Hoechst 33342. Cell staining was analyzed by flow cytometry.

### Immunoblotting

Cells were washed with PBS and lysed on ice with RIPA buffer (150 mM sodium chloride, 1.0% NP-40, 0.5% sodium deoxycholate, 0.1% sodium dodecyl sulfate, and 50 mM Tris, pH 8.0) containing protease inhibitor (Roche) for 30 min. The extracts were centrifuged at 13000 rpm for 15 min to remove the debris. The protein concentrations of the supernatants were determined using Quick Start™ Bradford 1x dye reagent. Equal amounts of protein were boiled with Laemmli buffer at 100⁰C for 5 min. The samples were briefly centrifuged and then separated on precast 4–20% gradient gels. The proteins were transferred to PVDF membranes, which were blocked with 5% skim milk at room temperature for one hour with gentle shaking. The blots were incubated with primary mouse SSRP1 or SPT16 antibodies at 4⁰C overnight. After the incubation, the blots were washed and incubated with HRP-conjugated, anti-mouse secondary antibodies for 1 hr at room temperature. As a control, the blots were incubated with HRP-conjugated mouse β-actin antibody at room temperature for 1 hr. The blots were washed for 1 min with Western Lightning Plus- ECL, and images were acquired using the BioRad ChemiDoc™ Touch Imaging System.

### Caspase-3 assay

The caspase-3 fluorogenic substrate Ac-DEVD-AMC was reconstituted in DMSO at a concentration of 1 mg/mL. Cells were lysed with 1X CCLR buffer, and the protein concentration was estimated using the Pierce BCA assay. The protein was used at a concentration of 2 μg/μL (50 μL). Ac-DEVD-AMC (20 μM in 20 mM HEPES, 10% glycerol, and 2 mM DTT) was added to each sample in a 96-well black plate (BD Falcon). The plates were incubated at 37⁰C for 1 hr, and then the fluorescence was measured.

### DNA minor groove binding

Cells (500,000) were treated with 4-OHT for five days and then serum-starved (0.5% FBS) for 48 hr. For the positive control, cells were treated with 1 μM of CBL0137 for 30 min. The cells were washed with PBS and stained with 3 μM Hoechst 33342 in 3% BSA-PBS at 37⁰C for 15 min.

### Flow cytometry

Flow cytometry was performed using LSRB II (Becton Dickinson). Histograms were constructed using WinList 3D software (Verity Software House).

### ATAC-seq

Two biological replicates were used for each condition. Cells were treated with 4-OHT for five days and then split. After attachment, the cells were cultured in medium containing 0.5% FBS for 48 hr. Cells (50,000) were prepared for ATAC-seq as previously described [35], with slight modifications. Briefly, the cells were washed with PBS, lysed, and incubated with the transposition reaction mixture at 37⁰C for 30 min. The DNA was purified using the Qiagen MinElute PCR Purification Kit and amplified using PCR primer 1 and barcoded primer 2 (1 cycle of 72°C for 5 min, 98°C for 30 sec, 13 cycles of 98°C for 10 sec, 63°C for 30 sec, 72°C for 1 min). The primers were custom synthesized by Integrated DNA Technologies (IDT) using previously described sequences [35]. The PCR products were purified using AmpureXP beads (Beckman Coulter). The resulting libraries were evaluated on High-Sensitivity D1000 ScreenTape using the TapeStation 4200 (Agilent Technologies). They were quantitated using the Kapa Biosystems qPCR quantitation kit for Illumina. The libraries were pooled, denatured, and diluted to 350 pM, with a 1% PhiX control library added. The resulting pool was loaded into a NovaSeq Reagent cartridge and sequenced using a NovaSeq6000 following the manufacturer’s recommended protocol (Illumina Inc.).

### RNA-seq

Two biological replicates were used for each condition. Cells were treated with 4-OHT for five days, then split and allowed to grow for 24 hr. The sequencing libraries were prepared from 500 ng total RNA using the mRNA HyperPrep kit (KAPA BIOSYSTEMS) following the manufacturer’s instructions. Briefly, PolyA RNA was purified using the mRNA Capture Beads. The purified RNA was fragmented and primed for cDNA synthesis. Fragmented RNA was reverse transcribed into first-strand cDNA using random primers. Second strand cDNA synthesis converted the cDNA:RNA hybrids to double-strand cDNA (dscDNA), with the 2nd strand marked by incorporating dUTP in place of dTTP. Pure Beads were used to separate the dscDNA from the second-strand reaction mix, resulting in blunt-ended cDNA. A single ‘A’ nucleotide was added to the 3’ ends of the blunt fragments. Multiple indexing adapters containing a single ‘T’ nucleotide on the 3’ end were ligated to the ends of the dscDNA to prepare them for hybridization onto a flow cell. The adapter-ligated libraries were amplified by PCR, purified using Pure Beads, and validated for the appropriate size using a 4200 TapeStation D1000 ScreenTape (Agilent Technologies, Inc.). The DNA libraries were quantitated using the KAPA Biosystems qPCR kit and pooled in an equimolar fashion. Each pool was denatured and diluted to 350 pM, with a 1% PhiX control library added. The resulting pools were loaded into NovaSeq Reagent cartridges for 100-cycle single-read analysis and sequenced with a NovaSeq6000 following the manufacturer’s recommended protocol (Illumina Inc.).

### Sample preparation for single-cell RNA-sequencing

Bone marrow and intestinal cells from vehicle- and tamoxifen-treated mice were used for single-cell gene expression using 10X Genomics technology. For bone marrow samples, mouse femurs were cleaned with alcohol, washed with ice-cold PBS containing antibiotics, and the metaphysis removed. The marrow was flushed with buffer (2 mM EDTA + 0.5% BSA in PBS) using a 23G syringe. The cells were gently mixed, filtered through a 40-μm cell strainer, and centrifuged at 300 g for 10 min. The pellets were resuspended in 0.04% BSA-PBS and counted using trypan blue. A total of 1 x 10^5^ cells were used for the analysis (1000 cells/μL).

For the intestinal samples, 2 cm of intestine above and below cecum, excluding cecum, were excised and flushed with ice-cold PBS containing gentamycin using a 20G needle. The intestine was placed in a dish with fresh PBS and cut open longitudinally. The walls were washed to remove any residual matter. The intestine was incubated in ice-cold PBS containing 10 mM EDTA (10 mL) at 4⁰C with gentle rocking for 30 min. The tissue was into ice-cold PBS and shaken with vigorous bursts to release the epithelial cells. The supernatant was centrifuged at 300 g for 5 min at 4⁰C. The pellet was washed once with ice-cold PBS, resuspended in PBS containing 1 mg/mL collagenase, and incubated at 37⁰C for 5 min. The tube was then placed immediately on ice, followed by the addition of 5% (v/v) FBS. The cell suspension was centrifuged, washed with PBS, and filtered through a 40-μm cell strainer. The cells were resuspended in 0.04 % BSA-PBS and counted using trypan blue. A total of 1 x 10^5^ cells were used for the analysis (1000 cells/μL).

### 10X Genomics

Single-cell libraries were generated using the 10X Genomics platform. Cell suspensions were assessed by trypan blue using the Countess FL automated cell counter (ThermoFisher) to determine the concentration, viability, and absence of clumps and debris that could interfere with single-cell capture. The cells were loaded into the Chromium Controller (10X Genomics), where they were partitioned into nanoliter-scale Gel Beads-in-emulsion with a single barcode per cell. Reverse transcription was performed, and the resulting cDNA was amplified. The full-length amplified cDNA was used to generate gene expression libraries by enzymatic fragmentation, end-repair, a-tailing, adapter ligation, and PCR to add Illumina-compatible sequencing adapters. The resulting libraries were evaluated on D1000 ScreenTape using the TapeStation 4200 (Agilent Technologies) and quantitated using the Kapa Biosystems qPCR quantitation kit for Illumina. The libraries were then pooled, denatured, and diluted to 300 pM, with 1% PhiX control library added. The resulting pool was loaded into the appropriate NovaSeq Reagent cartridge and sequenced on a NovaSeq6000, following the manufacturer’s recommended protocol (Illumina Inc.).

### Bioinformatics analysis

RNA-seq fastq files were aligned to the mouse mm10 genome using STAR [53] with GENCODE v25 gene annotation. Differential gene expression was assessed using DESeq2 [54] with default parameters. Genes with a 1.5-fold change between conditions and FDR < 0.05 were considered differentially expressed. GSEA was run using the MSigDB website (Broad Institute), and GO enrichment was run using g:Profiler [55] (Ensembl v104). The gene lists with FDR q-value ≤ 1e-4 were considered significant. The baseMean column from the DESeq2 differential expression result table was used to classify genes based on expression levels. A list of unexpressed genes was created from the genes with baseMean 0. The genes with baseMean > 0 were used for ranking based on expression levels. For this ranking, the baseMean values were divided by gene length and then ranked and divided into four quantiles.

ATAC-seq fastq files were aligned to the same genome using bowtie2 [56] with the following arguments: --very-sensitive --no-mixed --no-unal --no-discordant. Duplicates were filtered using the Picard MarkDuplicates function. The bigwig files with CPM-normalized profiles were obtained using deepTools [57] and parameters maxFragmentlength and minFragmentlength to account for the fragment sizes. Pile-ups were calculated using the computeMatrix function in the scale-regions or reference-point modes and visualized using the plotProfile and plotHeatmap functions from deepTools. The mean density reads at the gene bodies were calculated using the ComputeMatrix function from deepTools with the following parameters: scale-regions -o results.tab.gz --regionBodyLength 5000 -a 0 -b 0 --binSize 50 -- missingDataAsZero --skipZeros. The results.tab files for each expression category from two replicates were combined into one table. Bins 21–100 were used for the calculation of the mean density reads. These parameters were compared between conditions using the paired t- test.

Single-cell RNA-seq analysis: the raw sequencing data were processed using Cellranger (version 5.0.0) software (10x Genomics). The filtered gene-barcode matrices, which contained the barcodes with the Unique Molecular Identifier (UMI) counts that passed the cell detection algorithm, were used for further analysis. The Seurat single-cell data analysis R package was used for read normalization, dimension reduction, and cell clustering [58]. Cells with low RNA content (detected transcripts < 200), higher mitochondrial RNA content (> 20% of total reads), or very high RNA content (> 5000 detected transcripts, likely dublets) were filtered out from the analysis. The mitochondrial RNA content filter was not applied for the intestine data as some of the cells had very high mtRNA content, as expected [59]. Additionally, the cell cycle scores for the S and G2/M phases were determined using the CellCycleScoring method based on the mouse orthologous genes for the cc.genes in the Seurat package. The lists of the cell cycle genes from Buettner et al [60] were used to calculate the S and G2/M scores for each cell. The score was the average expression of the cell cycle genes subtracted by the aggregated expression of the control feature sets, which had the same number of genes and similar average gene expression value as the cell cycle genes.

The genes used for the S phase were Mcm4, Exo1, Slbp, Gmnn, Cdc45, Msh2, Mcm6, Rrm2, Pold3, Blm, Ubr7, Mcm5, Clspn, Hells, Nasp, Rpa2, Rad51ap1, Tyms, Rrm1, Rfc2, Prim1, Brip1, Usp1, Ung, Pola1, Mcm2, Fen1, Tipin, Pcna, Cdca7, Uhrf1, Casp8ap2, Cdc6, Dscc1, Wdr76, E2f8, Dtl, Ccne2, Atad2, Gins2, Chaf1b, and Pcna-ps2. The G2/M genes were Nuf2, Psrc1, Ncapd2, Ccnb2, Smc4, Lbr, Tacc3, Cenpa, Kif23, Cdca2, Anp32e, G2e3, Cdca3, Anln, Cenpe, Gas2l3, Tubb4b, Cenpf, Dlgap5, Hjurp, Cks1brt, Gtse1, Bub1, Birc5, Ube2c, Rangap1, Hmmr, Ect2, Tpx2, Ckap5, Cbx5, Nek2, Ttk, Cdca8, Nusap1, Ctcf, Cdc20, Cks2, Mki67, Tmpo, Ckap2l, Aurkb, Kif2c, Cdk1, Kif20b, Top2a, Aurka, Ckap2, Hmgb2, Cdc25c, Ndc80, and Kif11.

The normalized and scaled UMI counts were calculated using the SCTransform method and regressed against the cell cycle scores. Dimension reductions, including principal component analysis (PCA) and UMAP were carried out. Data clustering was identified using shared nearest neighbor (SNN)-based clustering on the first 30 principal components. Marker genes for each cluster were identified using the default parameters of the FindMarkers method in the Seurat package. For the bone marrow data, cell annotation was done by the singleR [61] R package using the Immgen database as a reference. For the small intestine, marker genes from the published single-cell data [29, 30] were used to annotate the cell types.

Venn diagrams were built using https://bioinformatics.psb.ugent.be/webtools/Venn/.

GSEA was run using MSigDB https://www.gsea-msigdb.org/gsea/msigdb/.

GO was run using https://biit.cs.ut.ee/gprofiler/gost.

### Statistical analyses

All experiments, except scRNA sequencing, were run with at least two biological replicates and included additional technical replicates (independent well, slides or fields of view). Animals were randomized between groups based on genotype, gender and age. No blinding was done for experiments with mice, except pathology examination of H&E stained slides. Mice experiments including measurement of mouse viability and weight included groups of 5 and more mice of the same gender. Other animal experiments including collection of samples for different assays were done with at least 2 animals of the same gender per group. Exact number of animals are provided in figure legends. Pathology examination was done without knowledge of mouse genotype. Student’s t-test was used to compare data between 2 groups. Details of the t-test used (paired or unpaired, one or two tails distribution are provided in figure legends. One-way ANOVA with Tukey posthoc tests was used to compare data between multiple groups.

## Acknowledgements

We are grateful for the help and support of the administrative office of the Cell Stress Biology Department of Roswell Park Comprehensive Cancer Center, specifically Mary Morgan, Sarah Tyrpak and Bruce Specht. We thank the members of the shared resources facilities for help and advice with the design and processing of the experiments, specifically Dr. Prashant Singh, Director of the Genomics Shared Resource, Dr. Joseph Spernyak, Director of the Translational Imaging Shared Resource, and Dr. Leslie Curtin, veterinarian in the Laboratory Animal Shared Resource. We also thank Shawn Matott, a senior systems analyst, for the help with the High Power Computer resources.

## Author contribution

IG performed part of the animal experiments and most of the cell-based experiments and drafted the manuscript. PS generated the mouse model, characterized the phenotypes of the mice upon FACT depletion, and edited the manuscript. AS generated the genetically-modified mouse embryonic stem cells and performed the *in vitro* fertilization. IT is an animal pathologist who reviewed and evaluated all H&E slides. AFS performed the EU incorporation assays and assisted with other experiments. MM and JW assisted with the bioinformatic analyses, and MM edited the manuscript. KG developed the concept of the study, analyzed all the data, and drafted and edited the manuscript.

## Availability of data and materials

The mouse model described in this study is available from MMRRC ID:46323. The datasets supporting the conclusions in this article are available in the GEO Datasets repository. scRNA-seq data are available as GSE189866, https://www.ncbi.nlm.nih.gov/geo/query/acc.cgi?acc=GSE189866; ATAC-seq and RNA-seq data are available as GSE189663, https://www.ncbi.nlm.nih.gov/geo/query/acc.cgi?acc=GSE189663; The datasets supporting the conclusions of this article are included as (Tables EV1–EV4).

## Expanded View Figure Legends

**Figure EV1.**
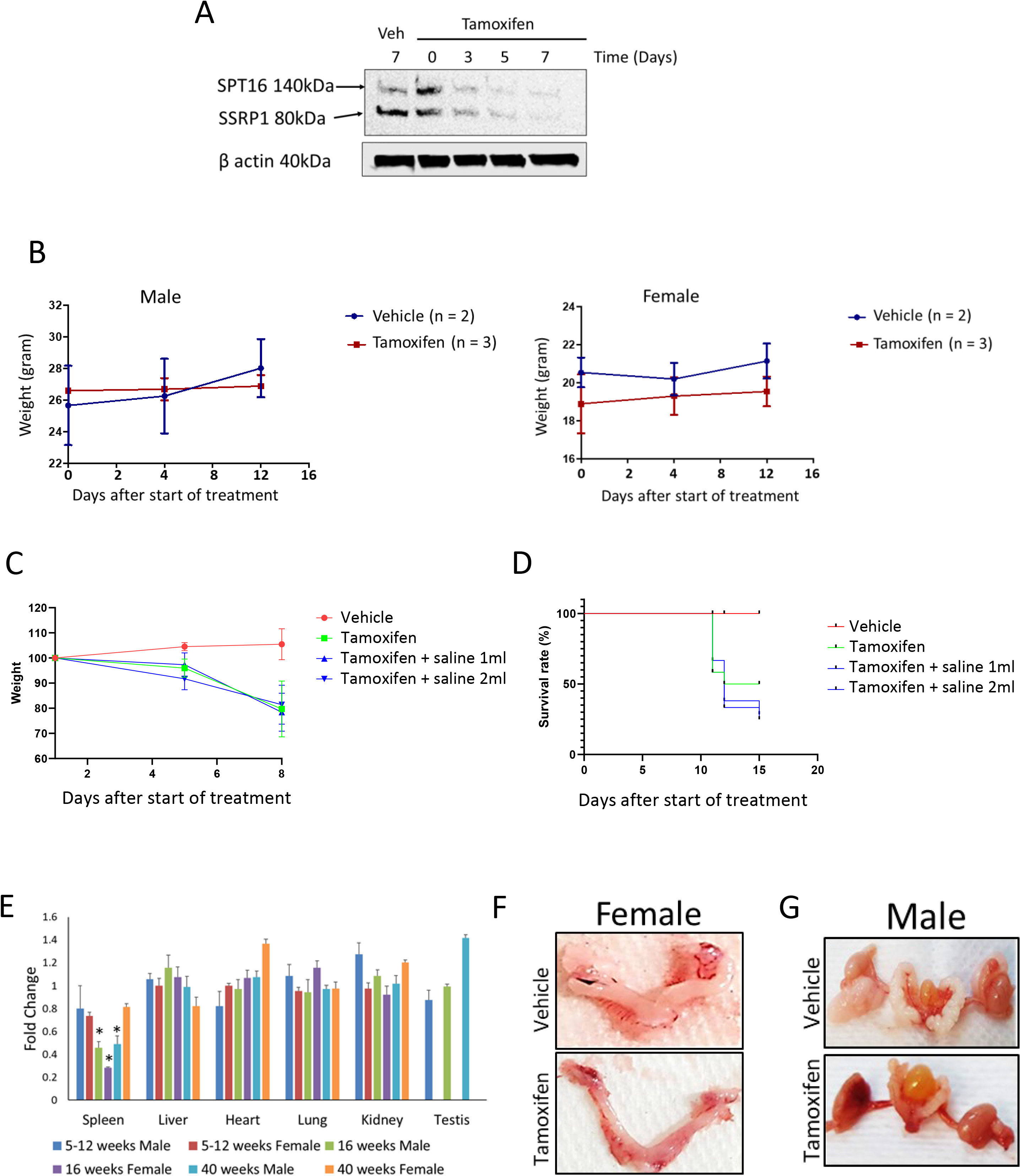
A. Kinetic of FACT protein subunit loss in spleen of *Ssrp1^fl/fl^; CreER^T2+/+^* mice upon tamoxifen administration. Western blotting of total protein extracts of spleen collected from mice at indicated time points. Veh is vehicle. B. Kinetic of weight changes of *Ssrp1^+/+^ ; CreER^T2+/+^* mice upon tamoxifen administration. Mean +/- SD. C. Effect of rehydration on weight of *Ssrp1^+/+^ ; CreER^T2+/+^* mice treated with vehicle or tamoxifen. Kinetic of weight loss in control and tamoxifen treated animals either provided with subcutaneous saline injection (1 or 2 ml per day) or not. Mean +/- SD, n = 3. D. Effect of rehydration on survival of *Ssrp1^+/+^ ; CreER^T2+/+^* mice treated with vehicle or tamoxifen. Kaplan-Mayer curves of control and tamoxifen treated groups either provided with subcutaneous saline injection (1 or 2 ml per day) or not. n = 3. E. Changes in organ weight in *Ssrp1^fl/fl^; CreER^T2+/+^* mice treated with tamoxifen versus vehicle treated animals. Mean +/- SD. * - p-value <0.05. n = 5. F – G. Representative photographs of female (F) and male (G) reproductive organs excised from vehicle and tamoxifen treated mice.

**Figure EV2.**
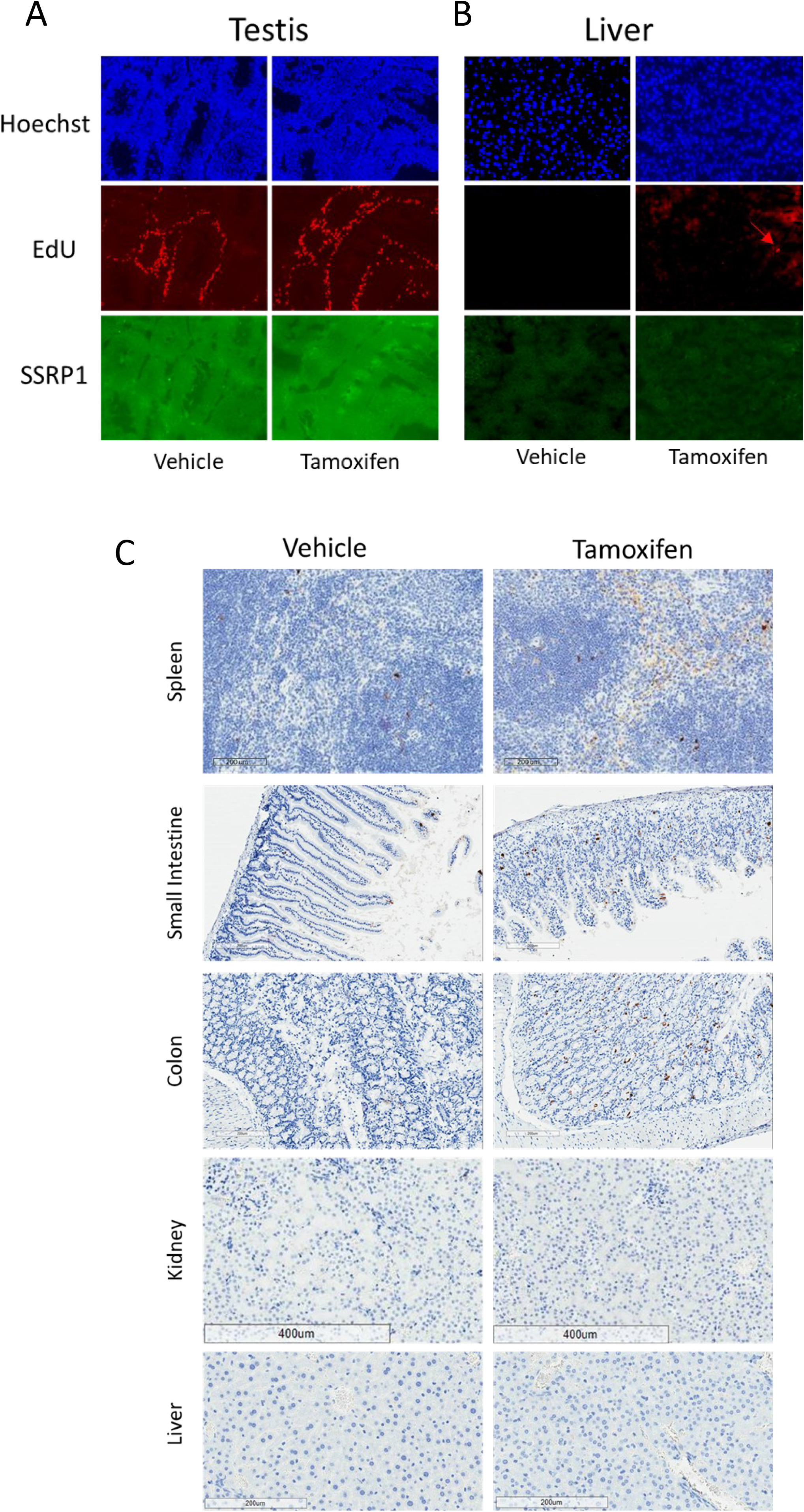
A-B. Effect of tamoxifen administration to *Ssrp1^fl/fl^; CreER^T2+/+^* mice on replication in testis (A) and liver (B). EdU was given to mice 3 days after stop of tamoxifen treatment for 1 hour followed by organ isolation and staining for total DNA (Hoechst), EdU and SSRP1. Red arrow indicates EdU positive nucleus in liver. C. Staining of sections of several organs of in *Ssrp1^fl/fl^; CreER^T2+/+^* mice for cleaved caspase 3 three days after stop of treatment with tamoxifen or vehicle.

**Figure EV3.**
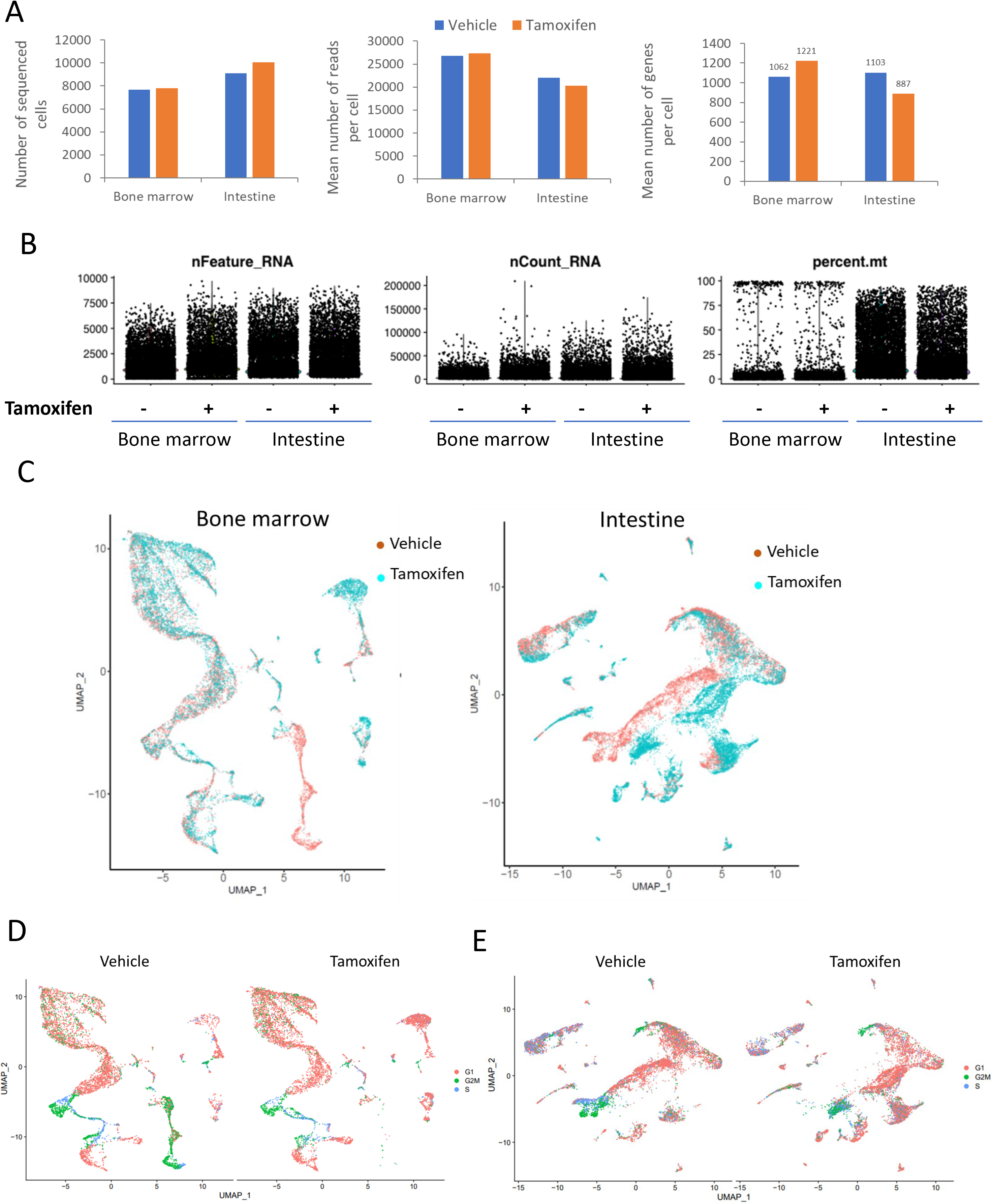
Analyses of scRNA sequencing of bone marrow and intestine of in *Ssrp1^fl/fl^; CreER^T2+/+^* mice next day after stop of treatment with vehicle or tamoxifen. A. Comparison of the numbers of sequenced cells, mean reads per cell and mean identified genes between samples. B. Dot plots showing the number of features (nFeature_RNA, the number of genes detected in each cell), RNA molecules (nCount_RNA, he total number of molecules detected within a cell) and proportion of reads corresponding to mitochondrial RNA (percent.mito) in each sample. C. Overlapped UMAP plots of bone marrow and intestinal samples color coded according to the treatment type. D-E. UMAP plots of bone marrow (D) and intestine (E) color-coded based on the expression of markers of cell cycle.

**Figure EV4.**
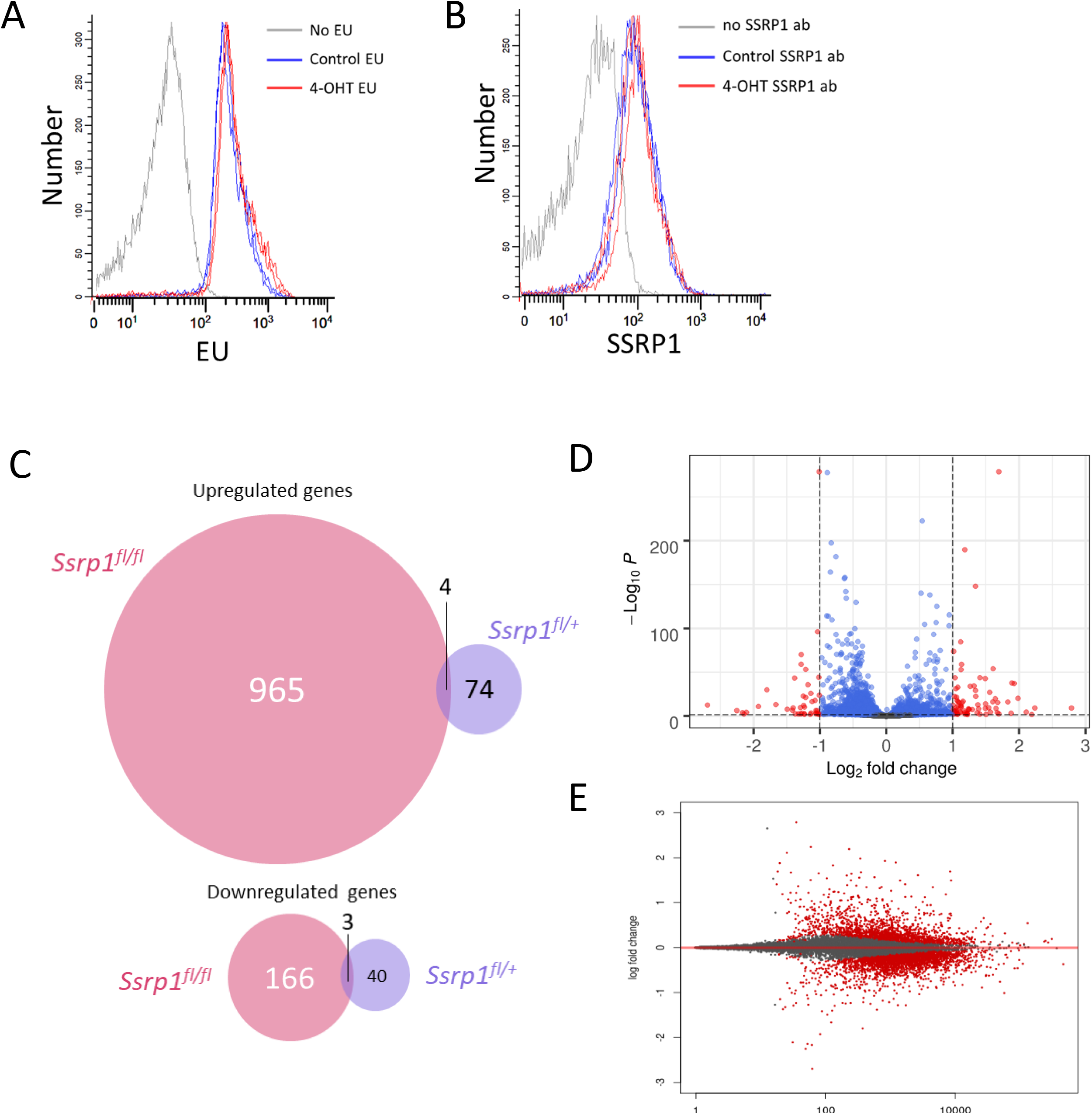
Comparison of changes in transcription in MSC of two genotypes treated with vehicle of 4-OHT for 5 days and then split and analyzed 24 hour after. A-B. EU incorporation assay (30 min) (A) and staining with SSRP1 antibody (B) of *Ssrp1^fl/+^; CreER^T2+/+^* MSC. Two replicates are shown. C - E. Data of RNA-seq experiment with two replicates of MSC. C. Venn diagrams showing number of up- and downregulated genes in homo- and heterozygous cells. D. Volcano plot of gene expression changes in heterozygous cells upon 4-OHT treatment. E. MA plot of gene expression changes in heterozygous cells upon 4-OHT treatment.

**Figure EV5.**
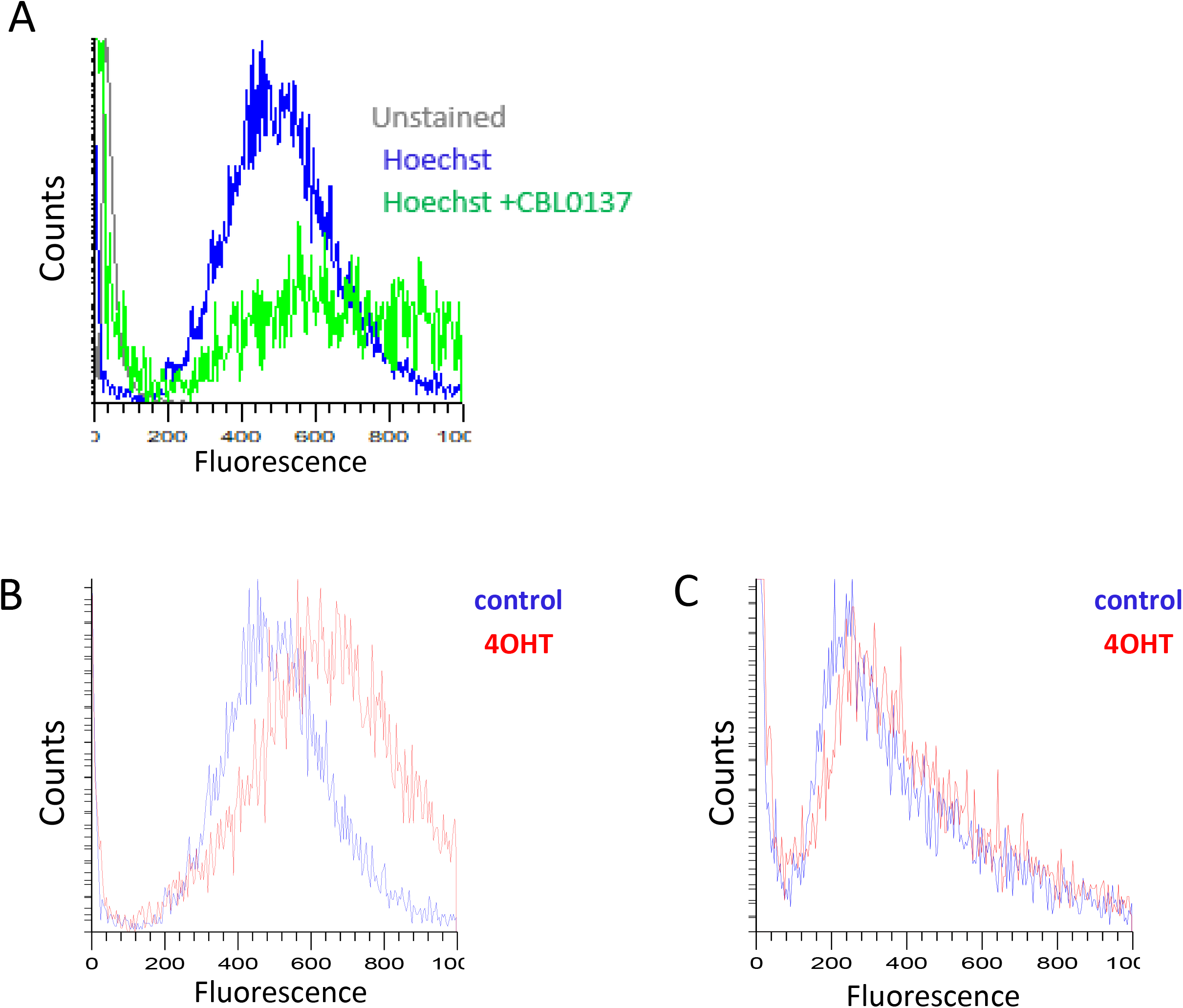
Assessment of general chromatin accessibility by staining DNA in live cells with Hoechst 33342. Histograms of fluorescent intensity of MSC stained with 3uM of Hoechst 33342 for 15 min. A. After incubation with CBL0137 (1uM) for 30 min. This plot also includes overlap of histogram of negative control (cells unstained with Hoechst). B. *Ssrp1^fl/fl^; CreER^T2+/+^* cells untreated or treated with 4OHT. C. *Ssrp1^fl/+^; CreER^T2+/-^* cells untreated or treated with 4OHT.

